# Associations between vascular risk factors and brain MRI indices in UK Biobank

**DOI:** 10.1101/511253

**Authors:** SR Cox, DM Lyall, SJ Ritchie, ME Bastin, MA Harris, CR Buchanan, C Fawns-Ritchie, MC Barbu, L de Nooij, LM Reus, C Alloza, X Shen, E Neilson, HL Alderson, S Hunter, DC Liewald, HC Whalley, AM McIntosh, SJ Lawrie, JP Pell, EM Tucker-Drob, JM Wardlaw, CR Gale, IJ Deary

**Author notes:** corresponding author: Simon R. Cox, Centre for Cognitive Ageing and Cognitive Epidemiology, Department of Psychology, The University of Edinburgh, Edinburgh EH8 9JZ, UK.

## Abstract

**Aims:** Several factors are known to increase risk for cerebrovascular disease and dementia, but there is limited evidence on associations between multiple vascular risk factors (VRFs) and detailed aspects of brain macro- and microstructure in large community-dwelling populations across middle- and older age.

**Methods and Results:** Associations between VRFs (smoking, hypertension, pulse pressure, diabetes, hypercholersterolaemia, BMI, and waist-hip ratio) and both global and regional brain structural and diffusion MRI markers were examined in UK Biobank (*N* = 9722, age range 44-77 years). A larger number of VRFs was associated with greater brain atrophy, lower grey matter volume, and poorer white matter health. Effect sizes were small (brain structural *R*^2^ ≤ 1.8%). Higher aggregate vascular risk was related to multiple regional MRI hallmarks associated with dementia risk: lower frontal and temporal cortical volumes, lower subcortical volumes, higher white matter hyperintensity volumes, and poorer white matter microstructure in association and thalamic pathways. Smoking pack years, hypertension and diabetes showed the most consistent associations across all brain measures. Hypercholesterolaemia was not uniquely associated with any MRI marker.

**Conclusion:** Higher levels of VRFs were associated with poorer brain health across grey and white matter macro- and microstructure. Effects are mainly additive, converging upon frontal and temporal cortex, subcortical structures, and specific classes of white matter fibres. Though effect sizes were small, these results emphasise the vulnerability of brain health to vascular factors even in relatively healthy middle and older age, and the potential to partly ameliorate cognitive decline by addressing these malleable risk factors.

## Introduction

With an increasingly ageing population, it is important to understand the neurobiological underpinnings of age-related cognitive impairment^1–3^. The functional sequelae of age-related cerebral decline carry a high financial, personal and societal burden, including impaired daily activities^4,5^, predict poorer health, and herald dementia, illness and death^6^. Dementia costs the UK more than £18 billion per year ^7^, and around 1% of global GDP in 2010^8^. The functional consequences of non-pathological brain ageing (which are much more prevalent than dementia) impose serious limitations on independence and quality of life in older age^9,10^. Efforts to understand the determinants of cerebral decline and quantify specific brain effects are urgently needed, especially with respect to modifiable factors which offer relatively direct pathways to intervention^1,8,10^.

Neurovascular health is an important correlate of preserved cognition in adult ageing^11,12^, yet significant gaps remain in our understanding of the links between vascular and cerebral ageing. Cerebral small vessel disease (CSVD; a constellation of clinical and imaging findings of presumed vascular aetiology^13^) causes ~45% of dementia and 20% of stroke worldwide^14^, though its pathophysiology and the interplay among its many possible determinants are not well understood^13^. Though the specific mechanisms by which these determinants, often known as vascular risk factors (VRFs), remain to be fully elucidated, anthropometric indices (waist:hip ratio and body mass index; WHR and BMI), blood glucose, elevated pulse pressure, chronic hypertension, diabetes and hypercholesterolaemia are all putative VRFs that have been associated with cerebrovascular complications^15–18^. The resultant damage to cerebral vasculature and increased vascular resistance are thought to deregulate cerebral blood flow, alongside blood brain barrier dysfunction, and could further lead to abnormal protein synthesis and formation of Alzheimer’s disease-typical plaques and tangles^15,18,19^.

In community-dwelling samples, the comparative importance of separate VRFs for the brain in relatively healthy ageing is unclear. Large scale comprehensive research designs are required to identify specifically which brain biomarkers are most sensitive to these potential effects, yet such data is scarce. Inconsistencies in the extant literature (discussed in ^20^, for example) may be partly down to low statistical power due to small sample sizes, and consideration of only one or few measures of risk and/or single brain MRI outcomes at any one time^20^. In non-pathological samples, effects are likely to be relatively subtle; well-powered, detailed MRI with multi-tissue analyses which can also account for multiple risk factors (and their tendency to cooccur) have been called for^21^.

UK Biobank represents one of the largest general population cohorts to have collected large-scale brain imaging data alongside information on VRFs among adults in middle and older age. This study examines total burden of vascular risk on global and regional measures of brain grey and white brain matter, derived from structural and diffusion MRI (dMRI) data in UK Biobank participants. We quantify the unique contributions to global and regional brain structure made by each simultaneously-modelled vascular risk factor. The wide age range further allows us to test the hypothesis that different VRFs may be more important for brain structure in midlife than in later life^22–24^.

## Methods

### Materials and procedure

When attending the assessment centre for an MRI scan, participants also provided demographic, health, and socioeconomic information in response to a series of touchscreen questions. To improve accuracy, they also took part in a nurse-led interview about their medical history, which included any self-reported diagnoses. (http://biobank.ctsu.ox.ac.uk/crystal/field.cgi?id=200). Participants were excluded from the present analysis if they reported having received a diagnosis of dementia, Parkinson’s disease or any other chronic degenerative neurological problem (including demyelinating diseases), brain cancer, brain haemorrhage, brain abscess, aneurysm, cerebral palsy, encephalitis, head injury, nervous system infection, head or neurological injury or trauma, stroke (N = 210). A total of 9,722 participants provided brain MRI scan data following exclusions, and automated and visual quality control (QC) by UK Biobank Imaging group. The present analyses were conducted under the approved UK Biobank research application number 10279.

#### Vascular Risk Factors

During medical history interview at the brain imaging appointment, participants also reported whether they had received a diagnosis of diabetes, hypertension, or hypercholesterolaemia. Data on cigarette smoking were also available from the touchscreen questionnaire. Blood pressure was collected twice, moments apart, using an Omron 705IT monitor. Pulse pressure was calculated as the log-transformed difference between mean systolic and mean diastolic pressure (or a single measure of each, where 2 were unavailable). Anthropometric measures were taken after participants had removed bulky clothing and shoes. Waist and hip measurements were conducted to provide WHR, and BMI was calculated as weight (kg) / height^2^ (m). For self-reported data, those who preferred not to answer or did not know were excluded from the analysis in all cases (<5%). Details of aggregate and latent VRF indices are described in Statistical Analysis.

#### MRI Acquisition and Processing

All brain MRI data was acquired on the same 3T Siemens Skyra scanner, according to a freely-available protocol (http://www.fmrib.ox.ac.uk/ukbiobank/protocol/V4_23092014.pdf), documentation (http://biobank.ctsu.ox.ac.uk/crystal/docs/brain_mri.pdf), and publication^25^. Briefly, the data were acquired with a standard Siemens 32-channel head coil. T_1_-weighted MPRAGE and T_2_-weighted FLAIR volumes were acquired in sagittal orientation at 1 × 1 × 1 mm and 1.05 × 1 × 1 mm resolution, respectively. The dMRI acquisition comprised a spin-echo echo-planar sequence with 10 T_2_-weighted (b≈0 s mm^−2^) baseline volumes, 50 b=1,000 s mm^−2^ and 50 b=2,000 s mm^−2^ diffusion weighted volumes, with 100 distinct diffusion-encoding directions and 2 mm isotropic voxels. The global tissue volumes, and white matter tract-averaged water molecular diffusion indices were processed by the UK Biobank team and made available to approved researchers as Imaging Derived Phenotypes (IDPs); the full details of the image processing and QC pipeline are available in an open access article^24^. These included total brain volume, grey matter volume, white matter hyperintensity (WMH) volume, subcortical volumes (accumbens, amygdala, caudate, hippocampus, pallidum, putamen, thalamus) and tract-averaged fractional anisotropy (FA) and mean diffusivity (MD) of the following white matter tracts: acoustic radiation, anterior thalamic, cingulum gyrus and parahippocampal, corticospinal, forceps major and minor, inferior fronto-occipital, inferior longitudinal, middle cerebellar peduncle, medial lemniscus, posterior thalamic, superior longitudinal, superior thalamic, uncinate. Extreme outlying data points (further than +/-4SDs from the mean were excluded case-wise (representing 0.001% of the total IDP datapoints analysed). Figure 1 shows the white matter tracts and subcortical sctructures.

**Figure 1.**
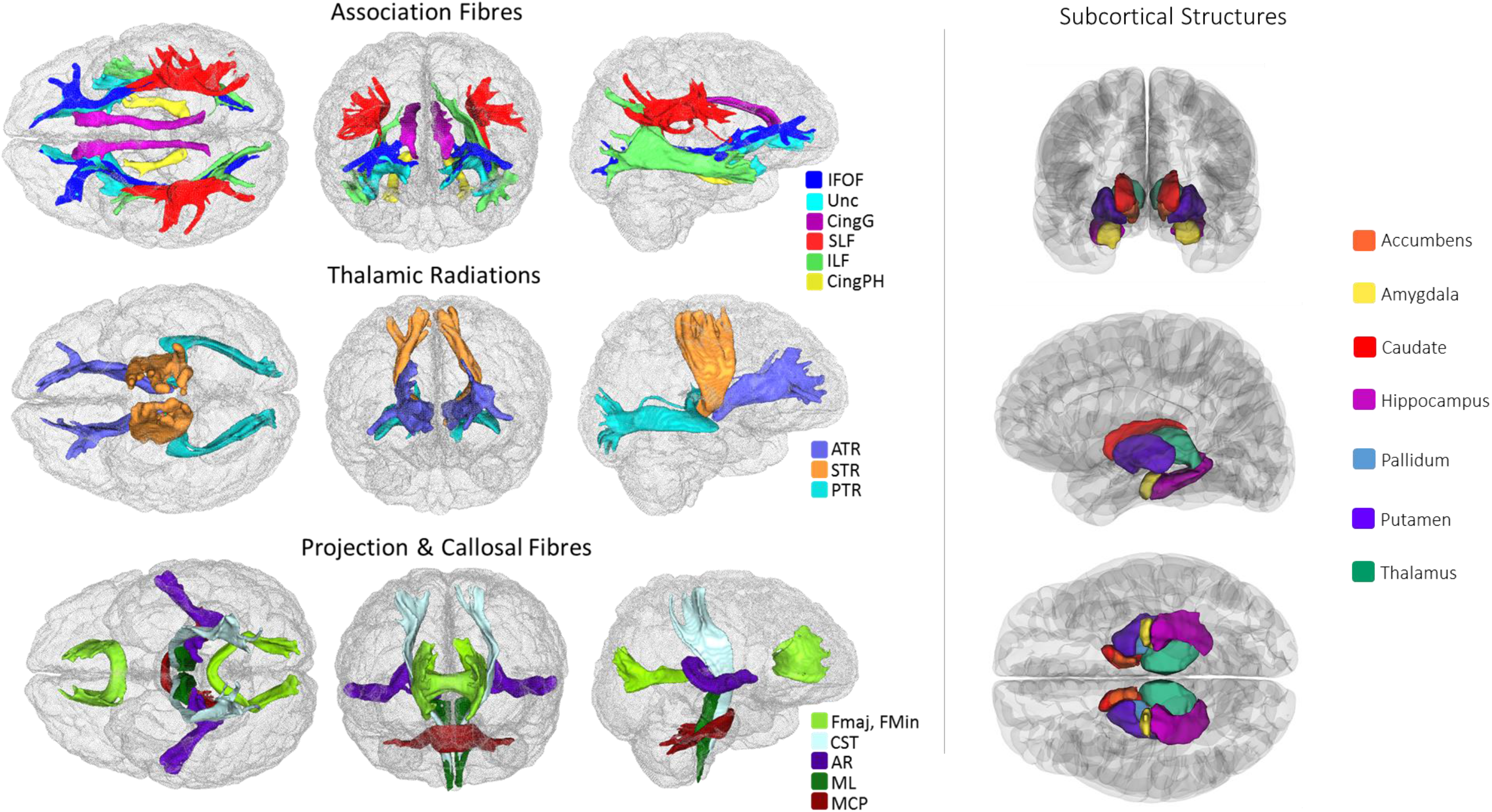
White matter tracts-of-interest (**left panel**) and subcortical structures (**right panel**) measured in the current study. IFOF = inferior fronto-occipital fasciculus, Unc = uncinate fasciculus, Cing = cingulum (gyrus and parahippocampal), SLF = superior longitudinal fasciculus, ILF = inferior longitudinal fasciculus, ATR = anterior thalamic radiation, STR = superior thalamic radiation, PTR = posterior thalamic radiation, Fmaj and Fmin (forceps major and minor), CST = corticospinal tract, AR = acoustic radiation, ML = medial lemniscus, MCP = middle cerebellar peduncle.

In addition to UK Biobank-provided IDPs, we conducted local processing and quality control of cortical reconstruction and segmentation, using FreeSurfer v5.3 on Ti-weighted volumes. Following visual inspection of the outputs (to check for aberrant surfaces and tissue segmentation failures, which were removed from analysis), a total of 8,975 participants had cortical surfaces available for analysis, 7,928 of whom also had casewise complete vascular, demographic and covariate data, and were used in the vertex-wise analyses. Surfaces were aligned vertex-wise into common space (the Freesurfer average template) and smoothed at 20mm full width at half maximum, allowing sample-wide analyses of volume across the cortical mantle.

### Analyses

All variables were visually inspected to ascertain whether they were distributed normally. BMI and WMH were log-transformed to correct a positively-skewed distribution. The ethnicity of the group was predominantly white, with 203 participants categorising themselves as non-white; this dichotomous variable for ethnicity was used as a covariate in all analyses. Pack years was calculated as the number of cigarettes per day divided by 20, and then multiplied by the number of years participants reported having smoked for. A latent measure of general white matter fractional anisotropy (*g*FA) and mean diffusivity (*g*MD) were derived using confirmatory factor analysis (‘cfa’ in the lavaan package) to index the high degree of covariance among white matter microstructural properties across the brain, as previously reported in this cohort^26^ and in others^27,28^. As described previously^26^, all tract measures (left and right) were entered separately into this analysis, correlated residuals between the left and right of each tract and between some other tracts were allowed, and based on low loadings (<0.3) of the Medial Lemniscus, Middle Cerebellar Peduncle and bilateral Parahippocampal Cingulum on the first factor, these measures were not included in the factor analysis. Tract loadings and model fit are shown in Supplementary Table 1.

We used two methods to capture the overall VRF load per individual. First, we derived an aggregate measure of vascular risk for each individual, counting instances of a self-reported diagnosis of hypertension, diabetes or hypercholesterolaemia, having ever smoked, having a BMI > 25^29,30^, and having a high WHR (>0.85 for females and >0.90 for males^31^). We also derived a latent factor of general vascular risk (*g*VRF) following prior work in an older cohort, using confirmatory factor analysis in structural equation modelling^32^. This latent measure captures, and depends upon, the tendency for vascular risk factors to co-occur. Using ‘cfa’ from the lavaan package, *g*VRF was derived from smoking pack years, diastolic and systolic blood pressure, BMI, WHR, self-reported hypertension, diabetes and hypercholesterolaemia. The model fit the data well, though loadings were inconsistent (range 0.175 to 0.758), with the factor more strongly loaded toward BMI and WHR (Supplementary Table 2).

First, we conducted descriptive analyses, testing associations between age and sex with each VRF (pack years, hypertension, pulse pressure, diabetes, hypercholesterolaemia, BMI and WHR) using linear regression (except for binary VRFs where logistic regression was used). We then examined associations between global MRI measures (total brain volume, grey matter volume, WMH, *g*FA and *g*MD) and overall and individual VRFs, before testing associations at the regional MRI level, which we now describe in more detail. We subsequently refer to total brain volume corrected for head size as global atrophy.

We ran linear regression analyses to quantify associations between overall load of VRFs (aggregate and latent VRF variables) and global MRI indices. We also included all age × VRF interactions; a significant interaction would mean that there was a difference association magnitude at different ages. In order to adequately correct our interaction term for the effects of sex, age, and age^2^ (i.e. non-linear), we also included sex × age and sex × age^2^ as covariate terms^33,34^. We also covaried for ethnicity, head size and differences in head positioning inside the scanner (X, Y and Z coordinates; http://biobank.ctsu.ox.ac.uk/showcase/label.cgi?id=110).

Next, we examined associations between each individual VRF and global MRI measures. Initially, we ran a separate simple model in which one VRF predicted each global MRI measure in turn, corrected for age, sex, ethnicity, head size, and scanner head position. We then re-ran these models to include a VRF × age interaction term, as described above. Finally, accounting for the fact that VRFs are positively correlated, we examined the unique contributions of each vascular risk factor (VRF) to global brain MRI measures by including all VRFs in one multiple linear regression for each MRI variable of interest. This approach allowed us to parse the relative contributions of each VRF – in the context of all others – to variance in brain MRI variables. To quantify the amount of variance in each brain imaging biomarker accounted for by VRFs, we compared the *R*^2^ of each model with that from a baseline model R^2^ in which the MRI measure was modelled with covariates only.

We then examined associations between regional MRI measures (white matter tract-specific FA and MD, vertex-wise cortical volume, and subcortical volumes) and the latent and aggregate measures of vascular risk. Finally, we examined the associations between individual VRFs and regional brain MRI. To do this, we first showed the basic associations for each individually-modelled VRF, before fitting a multiple regression for each of these regional MRI measures in which all individual VRFs were entered together. As before, all models also included age, sex, ethnicity, head size (for volumetric data) and head positioning confounds.

Statistical analyses were performed in R version 3.03 (https://www.r-project.org) except for cortical surface analyses which were performed using the SurfStat MATLAB toolbox (http://www.math.mcgill.ca/keith/surfstat) for Matrix Laboratory R2012a (The MathWorks, Inc, Natick, MA). We ensured that models showed acceptably low multicollinearity (variance inflation was ascertained using ‘vif’ in the ‘car’ package in R). Alpha was set at 0.05 for all analyses and results were corrected for multiple comparisons using FDR^35^ using ‘p.adjust’ package in R. Standardised coefficients are reported throughout to facilitate comparison of associations across all VRFs. Maps of the t-statistics for cortical analyses were displayed on the mantle such that negative associations with a VRF (i.e. lower volume) were always represented by the red end of the colour spectrum.

## Results

Participants were aged between 44.64 and 77.12 (M = 61.82, SD = 7.45) years, and participant characteristics are shown in Table 1. When modelling each VRF on age, sex and age × sex (Supplementary Table 3), females had significantly higher levels on all risk factors, and there were significant age × sex interactions for pack years (*β* = 0.039, *p* < 0.001), pulse pressure (*β* = −0.049, *p* < 0.001) and aggregate vascular risk (*β* = 0.025, *p* = 0.009). There was a significant association between greater vascular risk and older age across all VRFs (*β* range 0.083 to 0.498, *p* < 0.001) except for BMI which showed a weak relationship in the opposite direction (*β* = −0.027, *p* = 0.007). Plots of simple VRF trends with age are presented in Supplementary Figure 1. Consistent with the full UK Biobank cohort^36^, VRFs were generally modestly but significantly correlated (Supplementary Figure 2 and Supplementary Table 4). Older age was a relatively strong predictor of greater global atrophy and WMH volume, lower grey matter volume and *g*FA, and higher *g*MD (Supplementary Table 4; *β* range |0.254| to |0.586|, *p* < 0.001), consistent with prior reports from the initial release of UK Biobank MRI data (N ≈ 5000^26,37^). Sex differences in brain MRI measures were consistent with those previously-described in a smaller UK Biobank sample^38^.

**Table 1.**
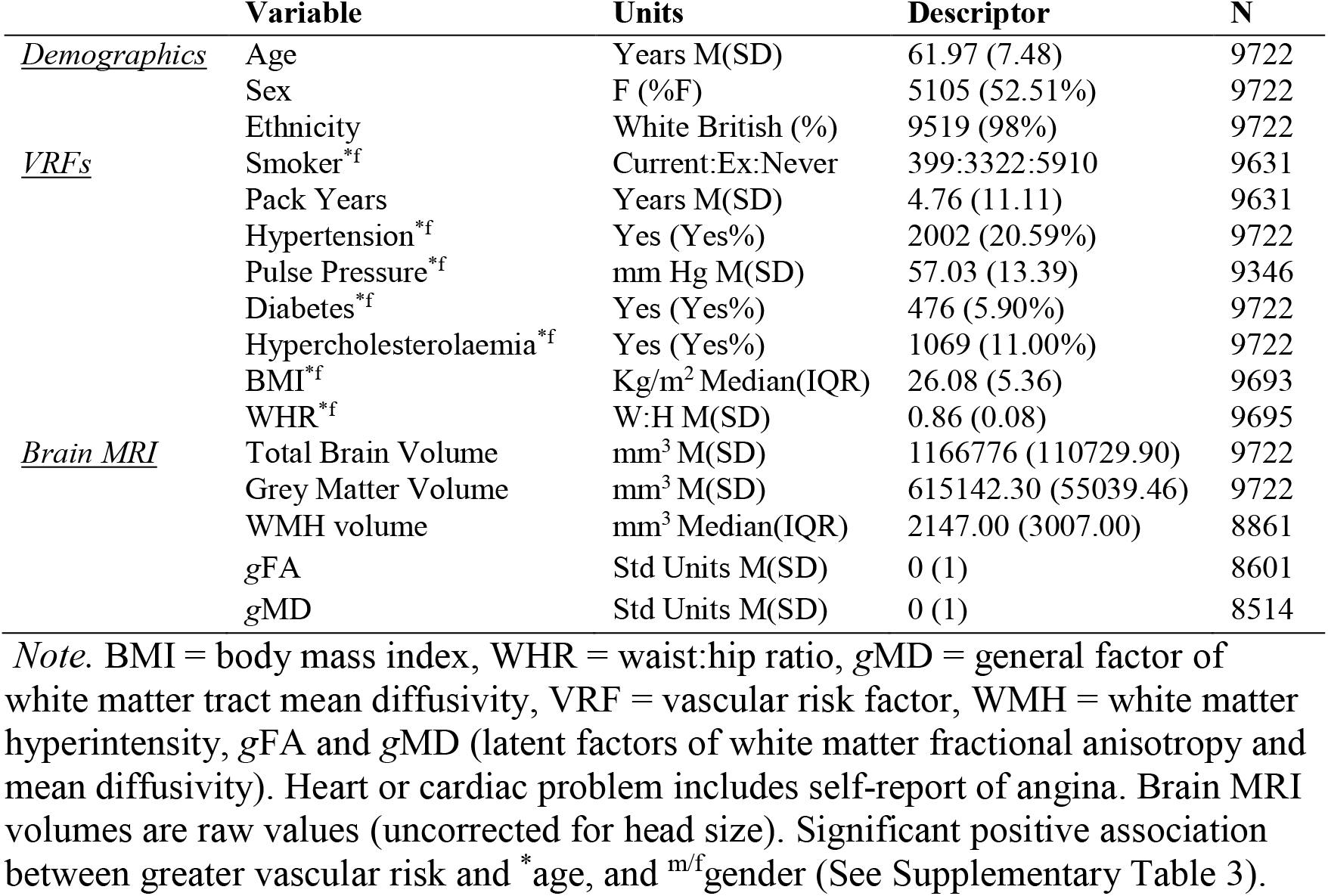
Participant characteristics.

### Global brain MRI analyses

#### General vascular risk

Associations between aggregate vascular risk and global MRI measures are reported in Table 2. Having a larger number of VRFs was associated with ostensibly ‘poorer’ global brain MRI health across all measures (*β* range |0.042| to |0.110|), accounting for ~1% of the variance in brain MRI measures beyond the contribution of covariates. Aside from the modest but significant positive interaction between age and aggregate VRF on higher *g*MD (interaction *β* = 0.036, *p* < 0.001; main effect *β* = 0.072, *p* < 0.001; indicating more VRFs are more strongly associated with less healthy white matter in older age), there was no evidence that associations between global brain measures and general vascular risk were stronger at different ages.

**Table 2.** Associations between individually- and simultaneously-modelled vascular risk factors on global brain MRI parameters.

| Vascular Risk Factors | Model | Global Atrophy |  | GM |  | WMH |  | gFA |  | gMD |  |
| --- | --- | --- | --- | --- | --- | --- | --- | --- | --- | --- | --- |
| | | $\beta$ | $p$ | $B$ | $p$ | $\beta$ | $p$ | $\beta$ | $p$ | $\beta$ | $p$ |
| **Aggregate VRF** | - | **-0.061** | **<0.001** | **-0.097** | **<0.001** | **0.110** | **<0.001** | **-0.042** | **<0.001** | **0.072** | **<0.001** |
| × age | - | 0.005 | 0.607 | -0.002 | 0.785 | -0.014 | 0.163 | -0.019 | 0.093 | **0.036** | **<0.001** |
| Additional R<sup>2</sup> |  | 0.005 |  | 0.008 |  | 0.010 |  | 0.002 |  | 0.007 |  |
| Total R<sup>2</sup> |  | 0.344 |  | 0.474 |  | 0.321 |  | 0.099 |  | 0.184 |  |
| Pack Years | *Single* | **-0.037** | **<0.001** | **-0.061** | **<0.001** | **0.076** | **<0.001** | **-0.036** | **<0.001** | **0.044** | **<0.001** |
|  | *Simultaneous* | **-0.029** | **<0.001** | **-0.046** | **<0.001** | **0.062** | **<0.001** | **-0.033** | **0.002** | **0.036** | **<0.001** |
| Hypertension | *Single* | **-0.027** | **0.001** | **-0.037** | **<0.001** | **0.097** | **<0.001** | **-0.084** | **<0.001** | **0.104** | **<0.001** |
|  | *Simultaneous* | **-0.022** | **0.012** | **-0.022** | **0.006** | **0.080** | **<0.001** | **-0.084** | **<0.001** | **0.094** | **<0.001** |
| Pulse Pressure | *Single* | **0.024** | **0.008** | 0.013 | 0.122 | **0.069** | **<0.001** | **-0.044** | **<0.001** | **0.074** | **<0.001** |
|  | *Simultaneous* | **0.033** | **<0.001** | **0.024** | **0.004** | **0.053** | **<0.001** | **-0.033** | **0.005** | **0.058** | **<0.001** |
| Diabetes | *Single* | **-0.044** | **<0.001** | **-0.066** | **<0.001** | **0.065** | **<0.001** | **-0.037** | **<0.001** | **0.045** | **<0.001** |
|  | *Simultaneous* | **-0.037** | **<0.001** | **-0.050** | **<0.001** | **0.038** | **<0.001** | -0.022 | 0.041 | **0.028** | **0.007** |
| High Cholesterol | *Single* | -0.003 | 0.712 | -0.006 | 0.434 | **0.023** | **0.009** | -0.007 | 0.520 | 0.016 | 0.099 |
|  | *Simultaneous* | 0.007 | 0.402 | 0.001 | 0.200 | 0.003 | 0.788 | 0.004 | 0.693 | 0.001 | 0.909 |
| BMI | *Single* | **-0.037** | **<0.001** | **-0.078** | **<0.001** | **0.053** | **<0.001** | 0.003 | 0.749 | 0.013 | 0.209 |
|  | *Simultaneous* | -0.012 | 0.240 | **-0.043** | **<0.001** | 0.003 | 0.790 | 0.026 | 0.041 | -0.028 | 0.022 |
| WHR | *Single* | **-0.060** | **<0.001** | **-0.101** | **<0.001** | **0.082** | **<0.001** | -0.000 | 0.981 | **0.041** | **0.002** |
|  | *Simultaneous* | **-0.036** | **0.007** | **-0.050** | **<0.001** | **0.048** | **<0.001** | 0.007 | 0.685 | 0.027 | 0.086 |
| VRF added R<sup>2</sup> |  | 0.004 |  | 0.011 |  | 0.018 |  | 0.011 |  | 0.015 |  |
| Total R<sup>2</sup> |  | 0.343 |  | 0.476 |  | 0.325 |  | 0.106 |  | 0.185 |  |

*Note.* Standardised betas ( $\beta$ ) and $p$ values are reported from regression models where vascular risk factors (VRFs) are regressed onto MRI measures, covarying for ethnicity, sex, age, sex × age and head position MRI confounds (volumetric data are also corrected for head size). Model denotes the results for each VRF modelled individually, and then when modelled simultaneously alongside all other VRFs. Additional R<sup>2</sup> refers to the amount of variance in MRI measures accounted for by the simultaneously-modelled VRFs, beyond covariates. Bold text denotes FDR q-value < 0.05. GM grey matter volume; BMI = body mass index; WHR = waist:hip ratio.

Alongside a measure of aggregate vascular risk, we quantified a latent factor of vascular risk (*g*VRF^32^), which showed a good fit to the data (Supplementary Figure 2, Supplementary Table 2). The measure of aggregate VRF and *g*VRF were strongly correlated (*r* = 0.795, *p* < 0.001). Whereas *g*VRF exhibited numerically larger association magnitudes with all global MRI measures apart from MD (Supplementary Table 5), these differences were modest; however, owing to the large N in the present study, we could detect that the *g*VRF associations were significantly larger for grey matter volume (*t*(9719) = 3.401, *p* <0.001) and WMH volume (*t*(8859) = 2.222, *p* = 0.026).

#### Individual vascular risk factors

Associations between global brain MRI measures and individual VRFs are also reported in Table 2. A greater number of pack years smoked, and a diagnosis of hypertension or diabetes were independently associated with putatively poorer global brain structural parameters (greater global atrophy, lower grey matter volume, more WMH, lower *g*FA and higher *g*MD; *β* range |0.022| to |0.104|). Higher BMI and WHR were both consistently associated with greater global atrophy, lower grey matter volume and higher WMH load, but both were non-significant for *g*FA, and only WHR was associated with higher *g*MD. Higher pulse pressure was associated with poorer white matter measures (higher WMH and MD, and lower FA), but was also related to less global atrophy (*β* = 0.24, *p* = 0.008) and was non-significant for grey matter volume. Finally, a diagnosis of hypercholesterolaemia was only significantly associated with greater WMH load, but with no other MRI index. No significant interactions were found between individual VRFs and global brain measures (Supplementary Table 6), and so age × VRF interactions were not further investigated at the brain regional level.

Simultaneous modelling of the associations between individual VRFs and general brain variables resulted in no, or only a minor attenuation of most effect sizes across measures (i.e. they were mostly independent contributions). Overall, magnitudes of associations between VRFs and global MRI variables were modest, explaining a small amount of additional variance beyond covariates. In order of smallest to largest, the incremental *R*^2^ explained by all VRFs was: global atrophy = 0.004, *g*FA = 0.011, grey matter volume = 0.011, *g*MD = 0.015, WMH = 0.018 (Table 2). The numerically largest associations were for hypertension on white matter measures (*β* range |0.080| to |0.094|, all *p* < 0.001), whereas self-reported hypercholesterolaemia did not make any significant unique contribution – above other VRFs – to any global MRI measure (*β* ≤ 0.007, *p* ≤ 0.198). The most notable effect of simultaneously modelling VRFs was that, whereas WHR still showed significant associations with global volumetric MRI measures (global atrophy, grey matter volume and WMH volume), the magnitude was attenuated by ~50%, and BMI was no longer significantly associated with any MRI measure except grey matter volume. This, however, indicates that variance in both BMI and WHR made unique contributions to lower grey matter volumes. Pack years, hypertension, pulse pressure and diabetes were still most consistently linked across all global brain outcomes.

### Regional brain MRI analyses

#### General vascular risk

We tested whether there was regional specificity underlying the global MRI associations between general and specific vascular risk. Vertex-wise cortical analysis revealed widespread significant associations between higher aggregate vascular risk and lower cortical volume (Figure 2). The FDR-corrected *q*-map illustrates the comparative sparing of dorsal motor/somatosensory and posterior cortical regions, while the t-map indicates largest effect sizes in the frontal and especially anterior lateral and medial temporal lobes.

**Figure 2.**
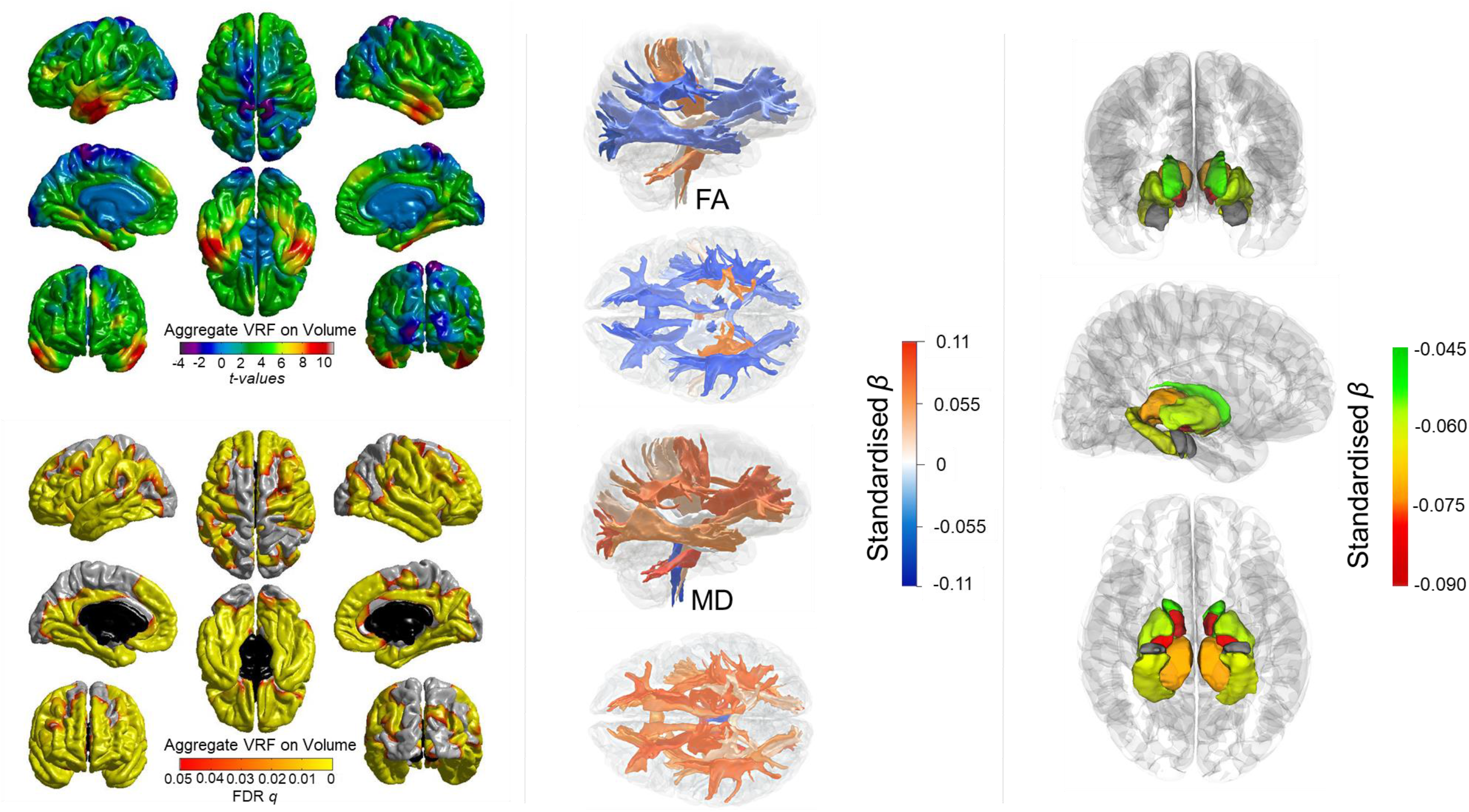
Associations between aggregate vascular risk and cortical volume (**left panel**), white matter tract-specific microstructure (**centre panel** showing right lateral and superior views), and subcortical volume (**right panel** showing, from top - bottom: anterior, lateral and inferior views). Higher aggregate vascular risk is associated with significantly lower cortical volume, lower FA and higher MD in the majority of white matter fibres, and lower subcortical volume, except for the amygdala (grey).

Results of the associations of aggregate vascular risk with white matter tract FA and MD, and with subcortical volumes are shown in Figure 2 and Supplementary Table 7. Higher aggregate vascular risk was associated with lower FA and higher MD and particularly implicated association and thalamic fibres. Unexpectedly, we also found some associations with projection fibres which were numerically among the largest magnitudes but in the opposite direction; higher aggregate risk was associated with *higher* FA in the corticospinal tract (*β* = 0.052), and middle cerebellar peduncle (*β* = 0.065), and also with *lower* MD in the medial lemniscus (*β* = −0.060). Higher aggregate vascular risk was also associated with generally lower subcortical volumes in all structures (*β* range −0.087 to −0.046, *p* < 0.001) except the amygdala (*β* = −0.006, *p* = 0.576).

The corresponding analysis for *g*VRF showed an almost identical pattern for cortical, white matter and subcortical measures (Supplementary Figure 3, Supplementary Table 8), though association magnitudes were, on average, slightly stronger throughout. For the vertex-wise analysis, this resulted in slightly higher t-values which were significant across slightly less restricted cortical loci, and was also the case for subcortical values.

#### Individual vascular risk factors

The patterning of significant associations of each individually-modelled VRF across the brain’s cortex are shown in Figure 3. All FDR-corrected (*q*) significant associations were in the expected direction (higher VRF with lower volume). Pulse pressure and hypercholesterolaemia showed no FDR-corrected significant associations with the cortex (Supplementary Figure 4). For pack years, hypertension, diabetes, BMI and WHR, a consistent pattern of associations emerged where the strongest effects in each case were in the lateral and medial temporal lobes. As with overall vascular risk, cortical associations toward the vertex were consistently absent. Medial and lateral frontal areas also showed significant associations, and relationships with occipital regions were most evident for smoking and diabetes.

**Figure 3.**
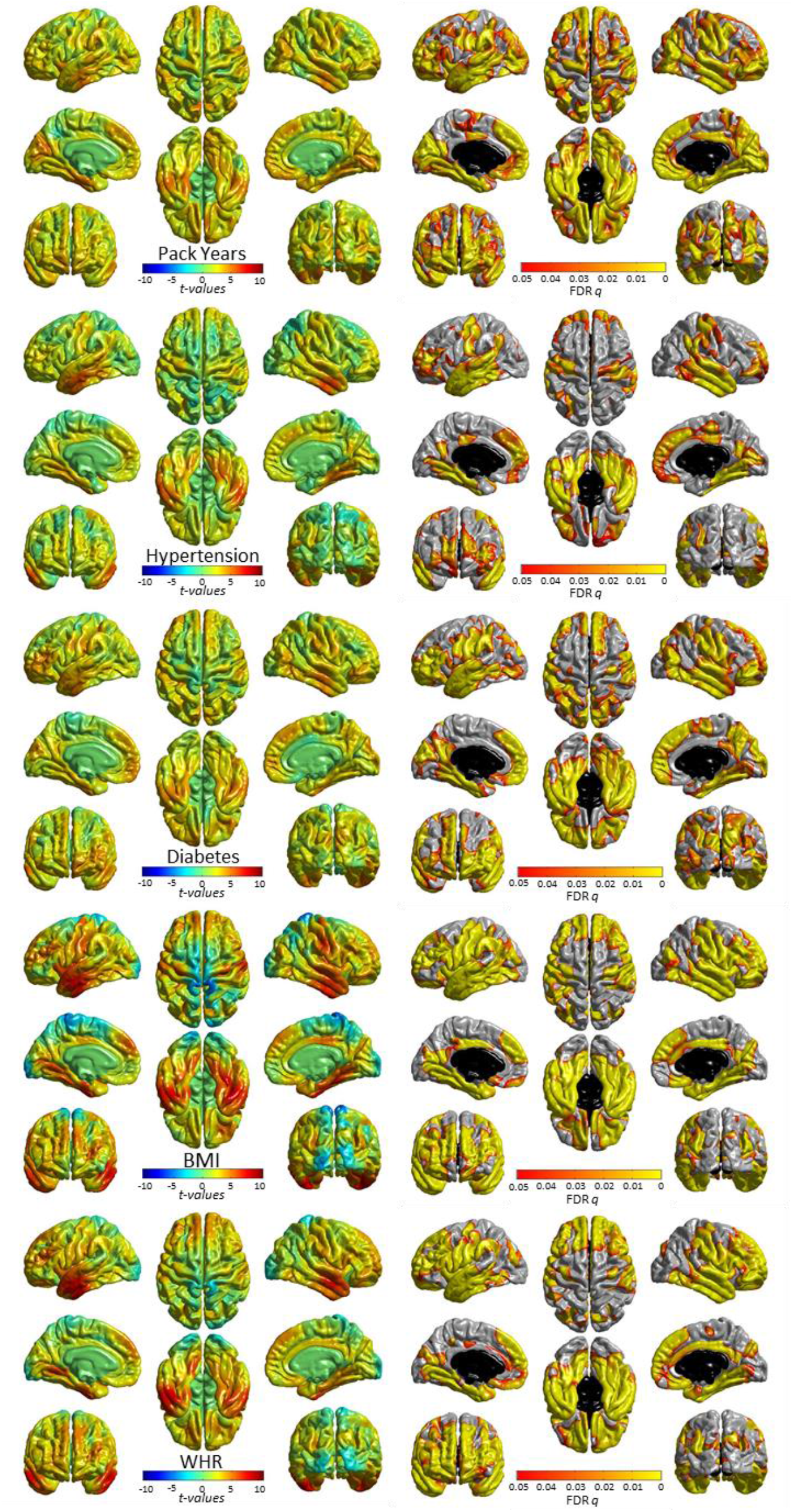
Significant associations (**left:** *t*-maps and **right:** FDR-corrected q-values) between cortical volume and vascular risk factors (modelled individually, alongside age, sex, ethnicity, head size and scanner head position confounds). See Supplementary Figure 3 for non-significant associations for pulse pressure and hypercholesterolaemia. T-maps are scaled with the same limits to aid comparison of relative effect size across risk factors.

Simultaneous modelling of individual VRFs across the cortex revealed the extent to which each VRF made a unique contribution to variance in regional volume, accounting for all other VRFs (Supplementary Figure 5). Though effect sizes were generally weaker, and the FDR-corrected loci were more restricted than when individually-modelled, the patterning of associations were largely unaltered. The common and unique patterns were more formally compared in the conjunction and conditional cortical analyses (Supplementary Figure 6). This emphasises clearly that i) individual VRFs make unique contributions to lower cortical volume at specific – common – foci: medial and anterior frontal, and temporal cortex, and ii) there were also regions which showed no overlap, indicating VRF-specific associations.

Associations between individually-modelled VRFs and white matter tract microstructure are reported in Figure 4 and Supplementary Tables 9 and 10. Thalamic and association fibres and the forceps minor showed the most consistent associations with lower FA and higher MD. These were driven by hypertension, pulse pressure, diabetes and pack years (*β* range |0.023| to |0.106|), in contrast to BMI and WHR whose associations were more consistent across projection bundles. Hypercholesterolaemia was not significantly associated with FA or MD in any tract.

**Figure 4.**
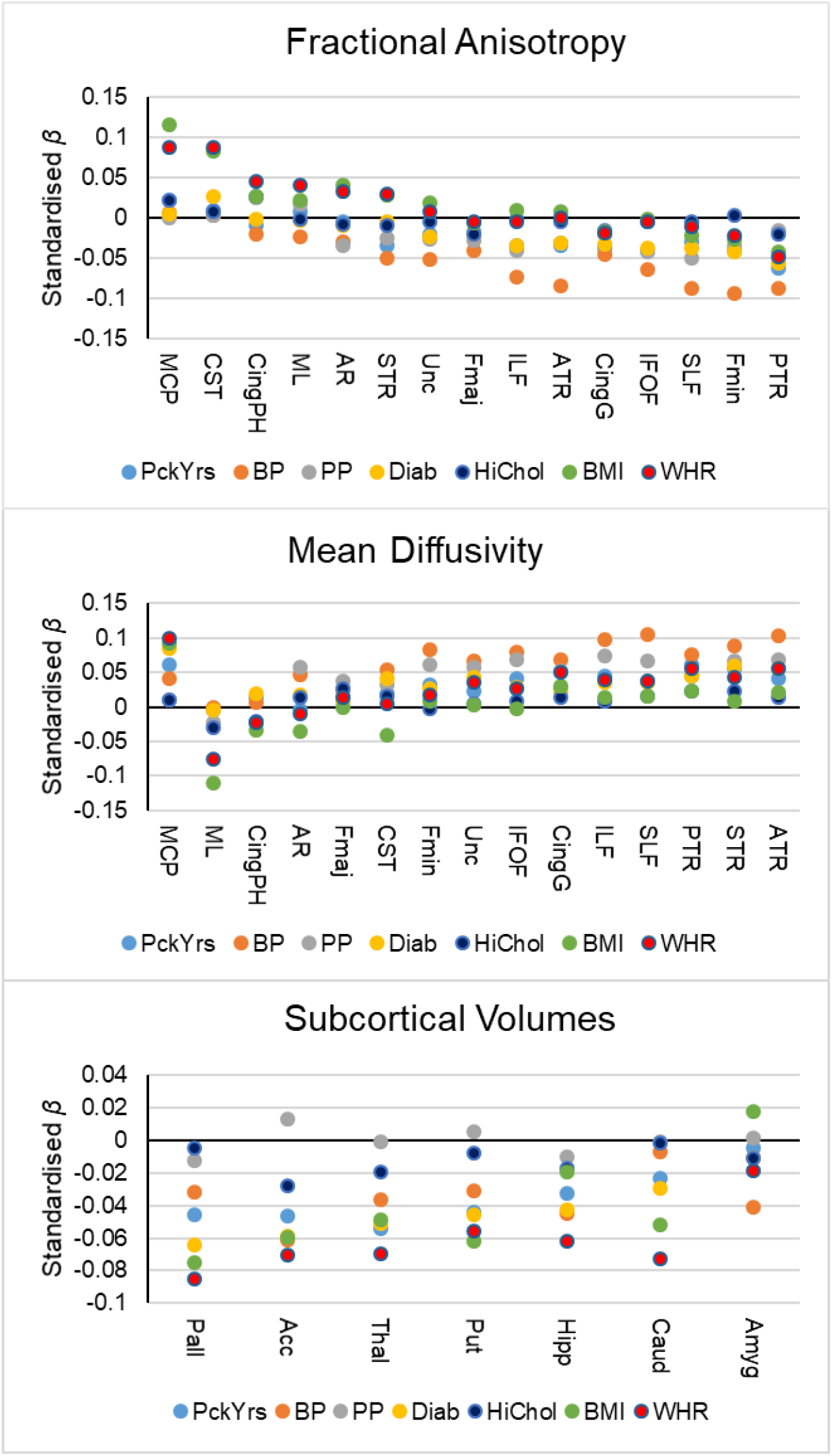
Standardised betas of associations between individually-modelled vascular risk factors and white matter tract FA (**top panel**), white matter tract MD (**centre panel**), and subcortical volumes (**bottom panel**). BP = hypertension, PP = pulse pressure, Diab = diabetes, HiChol = hypercholesterolaemia. AR = acoustic radiation, ATR = anterior thalamic radiation, Cing = cingulum (gyrus & parahippocampal), CST = corticospinal tract, Fmaj and Fmin (forceps major and minor), IFOF = inferior fronto-occipital fasciculus, ILF = inferior longitudinal fasciculus, MCP = middle cerebellar peduncle, ML = medial lemniscus, PTR = posterior thalamic radiation, SLF = superior longitudinal fasciculus, STR = superior thalamic radiation, Unc = uncinate fasciculus, Acc = accumbens, Amyg = amygdala, Caud = caudate, Hipp = hippocampus, Pall = pallidum, Put = putamen, Thal = thalamus.

Associations between subcortical volumes and VRFs are reported in Figure 4 and Supplementary Table 11. The majority of VRFs were significantly associated with lower volumes of all subcortical structures (*β* range |0.024| to |0.085|) except for the amygdala. However, hypercholesterolaemia was only associated with lower accumbens (*β* = −0.028, *p* = 0.002) and thalamus volumes (*β* = −0.019, *p* = 0.006), and pulse pressure showed no significant associations at all. Simultaneously modelling all VRFs for each tract (Supplementary Tables 11 and 12) and subcortical volume (Supplementary Table 13) did not substantially change this pattern of results, suggesting that effects are mainly independent from each other. However, high cholesterol was no longer significantly related to any subcortical volume.

## Discussion

### Interpretation

In this large, single-scanner sample of middle- and older-aged adults, associations between greater vascular risk and poorer brain health were small but significant across cortical, white matter, and subcortical tissue; there was a dose effect whereby the magnitude of association increased with the number of VRFs. Associations between vascular risk and brain structure did not differ appreciably across the sampled age range. We also provide insight into the relative contributions of different VRFs to brain health: greater pack years, a diagnosis of hypertension and diabetes each made unique contributions to poorer brain health across grey matter and WMH volume, white matter microstructure, and subcortical volume. Conversely, pulse pressure was mainly related to white but not grey matter measures, whereas WHR was only uniquely associated with volumetric grey and WMH (but not microstructure). A diagnosis of hypercholesterolaemia made no unique contributions to brain health beyond other risk factors. Throughout, effect sizes were small, accounting for less than 2% of the variance in brain structural measures.

Our results provide further evidence for regional cerebral vulnerability to VRFs in healthy individuals. On the cortex, associations between aggregate vascular risk and lower cortical volume shows strongest effects on frontal and especially anterior lateral and medial temporal lobes, rather than dorsal motor/somatosensory and posterior cortical regions. The cortical patterning in this large healthy sample agrees with loci associated with other markers of CSVD in community-dwelling adults^39,40^, and is strikingly consistent with the regional ischemic vulnerability of the brain to hypoperfusion in clinical samples^41^ and the pattern of atrophy common in ‘typical’ Alzheimer’s disease^42^.

We find that greater vascular risk is related to ‘poorer’ white matter microstructure (higher MD and lower FA) in specific classes of white matter tract. The association and thalamic radiations along with the forceps minor showed the most consistent significant relationships with vascular risk. These pathways also appear most susceptible to ageing, and we had previously hypothesised that this susceptibility and the phenomenon of age-related (statistical) de-differentiation might be partly driven by the disproportionately negative effects of environmental factors on these same fibres^25^. This also concurs with an explanation that these fibres connect the most metabolically active regions of the brain^43^ and may be at greatest risk of neurovascular ageing^15,19,41^. Finally, higher vascular risk was related to modestly lower subcortical volumes across the accumbens, caudate, hippocampus, pallidum, putamen and thalamus. Subcortical atrophy and WMH burden are also related to clinical diagnoses of dementia and vascular cognitive impairment^44^.

As far as we are aware, this is the largest single-scanner study of multiple VRFs and multi-modal structural brain imaging to-date. The high statistical power allowed us to reliably detect subtle and non-overlapping contributions of multiple individual VRFs to a large variety of brain health markers. Though there is some tendency for VRFs to co-occur, our simultaneous modelling indicated that smoking, hypertension, pulse pressure, diabetes and WHR each made unique statistical contributions to lower global brain and higher WMH volumes. Whereas it is possible that associations between obesity and brain structure are partly attributable to the findings that obesity promotes arterial stiffness, it is possible that the unique statistical contributions identified herein could pertain to other mechanisms through which body composition is linked to negative brain and cognitive endpoints, including metabolic and endocrine routes, which may have independent neurovascular consequences^17,24,45^. Similarly, the unique contributions to brain structure made by hypertension and smoking may indicate that the deleterious effects of smoking on the brain extend beyond putative alterations in hypertension^46^. This adds to the literature on the complex interplay between multiple sources of vascular risk and their associations with brain health.

### Limitations

The age range does not cover older ages (upper age limit was 77 years), restricting the degree to which our findings can be generalised to other populations. This may also have limited our scope to identify age-VRF interactions; such effects may be driven by very small associations in individuals who are much older than participants included here^23^. The UK Biobank imaging sample shows a tendency to live in less deprived areas than other UK Biobank participants^47^, who are already range restricted compared to the general population^48^, which may limit generalisability. Nevertheless, it is striking that associations between VRFs and brain structure are detectable even in these relatively healthy individuals, and effects may be larger in a more population-representative sample. Moreover, these data are cross-sectional, and cannot speak to lifelong trends in vascular risk, trajectories of brain regional decline nor important aspects such as lead-lag effects. Though we made attempts to remove individuals with neurological or neurodegenerative disorders, it is not possible to ascertain the degree to which the results reported here are driven by individuals with nascent age-related clinical neurodegenerative conditions.

The VRFs themselves represent different levels of fidelity – hypertension, diabetes and hypercholesterolaemia were binary rather than continuous measures, and all were based on self-report (albeit via a medical interview with a nurse). As a consequence, it is entirely possible that individual variability on continuous measures (such as blood cholesterol, or HbA1c) would prove more informative for brain outcomes. However, this alone cannot explain the absence of effects of hypercholesterolaemia, given the results for diabetes and hypertension. Though the simultaneous modelling of individual VRFs did not fundamentally alter the pattern of simple main effect, the combination of dichotomous and continuous variables could mean that any attenuations of estimates (from individual models) could partly be driven by this limitation, and should be interpreted with appropriate caution. On a related note, efficacy, dosage and time since diagnosis are all pertinent factors not measured here that are likely to have a bearing on our analyses (for example, individuals with a diagnosis of hypercholesterolaemia could be either taking medication or adapting their lifestyle appropriately). There are also likely to be many other lifestyle factors that are important for brain health that are not measured here (and serological markers which may offer greater precision than diagnosis information) which may well interact with differences in genetic susceptibility to such factors (e.g. ^49,50^, and should undoubtedly be a priority for future study in large brain imaging samples. This is especially important given that the majority of brain structural variation remains unexplained by the specific factors examined in this report.

Though the imaging acquisition and variety of sequences acquired can be considered state-of-the-art, there remain a number of inferential limitations with the available technology. For example, we are keen to observe the limitations of dMRI and of assuming that higher FA and lower MD automatically relate to ‘poorer’ white matter microstructure or health; there are many microstructural and methodological factors (including myelination, axonal bore, crossing fibres) that can influence the measurements of water molecular diffusion in the brain, necessitating a more nuanced interpretation^51^. This is evident in the conflicting directions of association between projection fibres compared to callosal, thalamic and association tracts. Taking the example of the middle cerebellar peduncle (which showed positive FA *and* MD betas) – in complex fibre structures with multiple crossing pathways, one can envisage a situation where FA and MD are positively correlated when more degraded transecting fibres provide less interference to molecular diffusion along the principal direction of the tract, thus leading to increases in both the directional coherence and the overall magnitude of water molecular diffusion.

### Summary

Elevated vascular risk in this large group of community-dwelling adults from the general population, was related to poorer brain health. A larger accumulation of vascular risk factors increased the magnitude of the association. The patterning of these effects was most pronounced in areas linked to elevated stroke and hypoperfusion susceptibility, and typical Alzheimer’s disease atrophy, and in the white matter pathways that facilitate their connectivity. Smoking, diabetes, and hypertension showed the most consistent associations across global and regional brain measures.

## Acknowledgements

We thank the UK Biobank participants and the UK Biobank team for their work in collecting, processing and disseminating these data for analysis. This research was conducted, using the UK Biobank Resource under approved project 10279, in The University of Edinburgh Centre for Cognitive Ageing and Cognitive Epidemiology (CCACE) (http://www.ccace.ed.ac.uk), part of the cross-council Lifelong Health and Wellbeing Initiative (MR/K026992/1). Funding from the Biotechnology and Biological Sciences Research Council (BBSRC) and Medical Research Council (MRC) is gratefully acknowledged. SRC, MEB, JMW, IJD were supported by MRC grants MR/M013111/1 and MR/R024065/1. IJD is additionally supported by the Dementias Platform UK (MR/L015382/1), and he, SRC and SJR by the Age UK-funded Disconnected Mind project (http://www.disconnectedmind.ed.ac.uk). SRC, SJR, CRB, MEB, IJD and EMT-D were supported by a National Institutes of Health (NIH) research grant R01AG054628. EMT-D was also supported by National Institutes of Health (NIH) research grant RO1HD083613. EMT-D is a member of the Population Research Center at the University of Texas at Austin, which is supported by NIH center grant P2CHD042849. JMW was supported by the Scottish Imaging Network: A Platform for Scientific Excellence (SINAPSE) collaboration (http://www.sinapse.ac.uk). CF-R is supported by Dementias Platform UK (DPUK), funded through the MRC (MR/L023784/2).

