## Supplementary Material for "Associations between vascular risk factors and brain MRI indices in UK Biobank"

**Supplementary Tables**

*Supplementary Table 1.* Tract loadings on a latent factor of white matter fractional anisotropy (gFA) and mean diffusivity (gMD)

| **Tract** | **Side** | **Loading FA** | **Loading MD** |
|---|---|---|---|
| Acoustic Radiation | L | 0.603 | 0.460 |
| | R | 0.621 | 0.535 |
| Anterior Thalamic Radiation | L | 0.786 | 0.815 |
| | R | 0.773 | 0.811 |
| Cingulum (Gyrus) | L | 0.458 | 0.745 |
| | R | 0.494 | 0.744 |
| Corticospinal Tract | L | 0.492 | 0.604 |
| | R | 0.497 | 0.627 |
| Forceps | Maj | 0.532 | 0.352 |
| Forceps | Min | 0.819 | 0.805 |
| Inferior Longitudinal Fasciculus | L | 0.813 | 0.784 |
| | R | 0.851 | 0.860 |
| Inferior Fronto-Occipital | L | 0.847 | 0.837 |
| | R | 0.847 | 0.887 |
| Posterior Thalamic Radiation | L | 0.643 | 0.661 |
| | R | 0.638 | 0.721 |
| Superior Longitudinal Fasciculus | L | 0.831 | 0.923 |
| | R | 0.848 | 0.919 |
| Superior Thalamic Radiation | L | 0.602 | 0.867 |
| | R | 0.590 | 0.841 |
| Uncinate Fasciculus | L | 0.661 | 0.544 |
| | R | 0.686 | 0.691 |
| CFI | | 0.971 | 0.979 |
| TLI | | 0.959 | 0.958 |
| RMSEA | | 0.059 | 0.069 |
| SRMR | | 0.033 | 0.037 |

*Note.* Standardised loadings are provided. Bilateral measures were introduced separately. CFI: Comparative Fit Index, TLI: Tucker-Lewis Index, RMSEA: Root Mean Squared Error of Approximation, SRMR: Standardised Root Mean Square Residual.

*Supplementary Table 2.* Loadings and model fit for a factor analysis of vascular risk.

| VRF | Loading |
|---|---|
| Pack Years | 0.237 |
| Hypertension | 0.313 |
| SBP | 0.320 |
| DBP | 0.255 |
| Diabetes | 0.307 |
| Hypercholesterolaemia | 0.175 |
| BMI | 0.588 |
| WHR | 0.758 |
| CFI | 0.981 |
| TLI | 0.957 |
| RMSEA | 0.041 |
| SRMR | 0.022 |

*Note.* Standardised loadings are provided. CFI: Comparative Fit Index, TLI: Tucker-Lewis Index, RMSEA: Root Mean Squared Error of Approximation, SRMR: Standardised Root Mean Square Residual. See also Supplementary Figure 5.

*Supplementary Table 3.* Associations between vascular risk factors, age and sex.

| Variable | Age | Sex | Age × Sex |
|---|---|---|---|
| Pack Years | 0.083 (<0.001) | 0.080 (<0.001) | 0.039 (<0.001) |
| Hypertension† | 0.464 (<0.001) | 0.218 (<0.001) | -0.032 (0.251) |
| Pulse Pressure | 0.385 (<0.001) | 0.113 (<0.001) | -0.049 (<0.001) |
| Diabetes† | 0.356 (<0.001) | 0.360 (<0.001) | -0.036 (0.489) |
| Hypercholesterolaemia† | 0.498 (<0.001) | 0.224 (<0.001) | -0.071 (0.050) |
| BMI | -0.027 (0.007) | 0.121 (<0.001) | -0.012 (0.150) |
| WHR | 0.083 (<0.001) | 0.666 (<0.001) | 0.002 (0.773) |
| *g*VRF | 0.126 (<0.001) | 0.529 (<0.001) | 0.001 (0.932) |
| VRFtot | 0.166 (<0.001) | 0.283 (<0.001) | 0.025 (0.009) |

*Note.* Standardised $\beta$ (*p*-values) are reported except †logistic regression (predictors were scaled). BMI = body mass index, WHR = waist:hip ratio, *g*VRF = general factor of vascular risk, VRFtot = aggregate measure of vascular risk. Female coded as 0.

*Supplementary Table 4.* Associations among study variables.

| | Age | *g*VRF | VRFtot | PackYrs | HiBP | PP | Diab | HiChol | BMI[a] | WHR | Atrophy | GM | *g*FA | *g*MD | WMH[a] |
|---|---|---|---|---|---|---|---|---|---|---|---|---|---|---|---|
| **Age** | ***** | 0.174 | 0.193 | 0.092 | 0.181 | 0.392 | 0.077 | 0.146 | -0.017 | 0.143 | -0.547 | -0.586 | -0.254 | 0.363 | 0.394 |
| ***g*VRF** | <0.001 | ***** | 0.795 | 0.282 | 0.373 | 0.278 | 0.365 | 0.209 | 0.700 | 0.902 | -0.231 | -0.355 | -0.007 | 0.121 | -0.027 |
| **VRFtot** | <0.001 | <0.001 | ***** | 0.361 | 0.518 | 0.174 | 0.354 | 0.390 | 0.587 | 0.616 | -0.213 | -0.295 | -0.053 | 0.135 | 0.077 |
| **PackYrs** | <0.001 | <0.001 | <0.001 | ***** | 0.080 | 0.054 | 0.102 | 0.055 | 0.136 | 0.176 | -0.109 | -0.147 | -0.047 | 0.075 | 0.083 |
| **HiBP** | <0.001 | <0.001 | <0.001 | <0.001 | ***** | 0.195 | 0.152 | 0.163 | 0.183 | 0.176 | -0.145 | -0.173 | -0.115 | 0.163 | 0.137 |
| **PP** | <0.001 | <0.001 | <0.001 | <0.001 | <0.001 | ***** | 0.087 | 0.051 | 0.072 | 0.166 | -0.213 | -0.247 | -0.112 | 0.219 | 0.180 |
| **Diab** | <0.001 | <0.001 | <0.001 | <0.001 | <0.001 | <0.001 | ***** | 0.102 | 0.173 | 0.175 | -0.103 | -0.137 | -0.047 | 0.072 | 0.072 |
| **HiChol** | <0.001 | <0.001 | <0.001 | <0.001 | <0.001 | <0.001 | <0.001 | ***** | 0.094 | 0.125 | -0.087 | -0.114 | -0.049 | 0.059 | 0.055 |
| **BMI[a]** | 0.101 | <0.001 | <0.001 | <0.001 | <0.001 | <0.001 | <0.001 | <0.001 | ***** | 0.447 | -0.069 | -0.117 | 0.031 | 0.008 | 0.015 |
| **WHR** | <0.001 | <0.001 | <0.001 | <0.001 | <0.001 | <0.001 | <0.001 | <0.001 | <0.001 | ***** | -0.212 | -0.348 | 0.035 | 0.081 | -0.133 |
| **Atrophy** | <0.001 | <0.001 | <0.001 | <0.001 | <0.001 | <0.001 | <0.001 | <0.001 | <0.001 | <0.001 | ***** | 0.850 | 0.159 | -0.271 | -0.127 |
| **GM** | <0.001 | <0.001 | <0.001 | <0.001 | <0.001 | <0.001 | <0.001 | <0.001 | <0.001 | <0.001 | <0.001 | ***** | 0.181 | -0.291 | -0.098 |
| ***g*FA** | <0.001 | 0.538 | <0.001 | <0.001 | <0.001 | <0.001 | <0.001 | <0.001 | 0.004 | 0.001 | <0.001 | <0.001 | ***** | -0.744 | -0.394 |
| ***g*MD** | <0.001 | <0.001 | <0.001 | <0.001 | <0.001 | <0.001 | <0.001 | <0.001 | 0.470 | <0.001 | <0.001 | <0.001 | <0.001 | ***** | 0.343 |
| **WMH[a]** | <0.001 | 0.011 | <0.001 | <0.001 | <0.001 | <0.001 | <0.001 | <0.001 | 0.171 | <0.001 | <0.001 | <0.001 | <0.001 | <0.001 | ***** |

*Note.* Pearson's *r* (above diagonal) and *p* value (below diagonal) reported. [a] log transformed. HiBP = self-report of hypertension, HiChol = self-reported hypercholesterolaemia, BMI = body mass index, WHR = waist:hip ratio, GM = grey matter volume corrected for head size, WMH = white matter hyperintensity volume corrected for head size, *g*FA = latent measure of general white matter mean diffusivity *g*MD = latent measure of general white matter mean diffusivity, *g*VRF = latent measure of general vascular risk, VRFtot = aggregate measure of vascular risk.

*Supplementary Table 5.* Associations of general vascular risk indices, and their age interactions, on global brain MRI.

| | **Global Atrophy** $\beta$ (*p*-value) | **GM** $\beta$ (*p*-value) | **WMH** $\beta$ (*p*-value) | ***g*FA** $\beta$ (*p*-value) | ***g*MD** $\beta$ (*p*-value) |
|---|---|---|---|---|---|
| ***Latent* VRF** | **-0.067 (<0.001)** | **-0.119 (<0.001)** | **0.125 (<0.001)** | **-0.036 (0.004)** | **0.079 (<0.001)** |
| × age | 0.002 (0.866) | -0.012 (0.161) | -0.010 (0.339) | -0.012 (0.324) | **0.033 (0.006)** |
| Additional $R^2$ | 0.003 | 0.010 | 0.010 | 0.001 | 0.005 |
| Total $R^2$ | 0.344 | 0.476 | 0.322 | 0.097 | 0.182 |
| ***Aggregate* VRF** | **-0.061 (<0.001)** | **-0.097 (<0.001)** | **0.110 (<0.001)** | **-0.042 (<0.001)** | **0.072 (<0.001)** |
| × age | 0.005 (0.607) | -0.002 (0.785) | -0.014 (0.163) | -0.019 (0.093) | **0.036 (<0.001)** |
| Additional $R^2$ | 0.005 | 0.008 | 0.010 | 0.002 | 0.007 |
| Total $R^2$ | 0.344 | 0.474 | 0.321 | 0.099 | 0.184 |

*Note.* Standardised betas ($\beta$) and *p* values reported from regression models where latent or aggregate vascular risk are regressed onto MRI measures, covarying for ethnicity, sex, age, sex × age and head position MRI confounds (volumetric data are also corrected for head size). GM: grey matter volume corrected for head size, *g*FA: general factor of fractional anisotropy, *g*MD: general factor of mean diffusivity, WMH: white matter hyperintensity volume, VRF: vascular risk factor. Additional R2 refers to the amount of variance in MRI measures accounted for by the simultaneously-modelled VRFs, beyond covariates. Bold type denotes FDR q-value < 0.05. Aggregate results are also presented in Table 2, but are shown here for comparison with latent VRF results.

*Supplementary Table 6.* Associations between individually-modelled vascular risk factors and their age interactions, on brain MRI parameters.

| **Vascular Risk Factors** | **Global Atrophy** *β* | *p* | **GM** *B* | *p* | **WMH** *β* | *p* | ***g*FA** *β* | *p* | ***g*MD** *β* | *p* |
|---|---|---|---|---|---|---|---|---|---|---|
| Pack Years × age | 0.012 | 0.207 | 0.013 | 0.122 | -0.008 | 0.453 | -0.012 | 0.295 | 0.025 | 0.030 |
| Hypertension × age | 0.020 | 0.033 | 0.011 | 0.164 | -0.002 | 0.856 | -0.018 | 0.117 | 0.023 | 0.037 |
| Pulse Pressure × age | -0.001 | 0.911 | 0.004 | 0.679 | -0.005 | 0.615 | -0.004 | 0.742 | -0.008 | 0.527 |
| Diabetes× age | 0.008 | 0.383 | 0.004 | 0.598 | -0.001 | 0.885 | -0.002 | 0.890 | 0.017 | 0.124 |
| HiCholesterol × age | -0.009 | 0.364 | -0.015 | 0.086 | -0.023 | 0.020 | 0.015 | 0.202 | -0.007 | 0.500 |
| BMI × age | 0.009 | 0.296 | -0.002 | 0.829 | -0.012 | 0.169 | 0.004 | 0.735 | 0.003 | 0.766 |
| WHR × age | -0.012 | 0.291 | -0.025 | 0.015 | -0.021 | 0.089 | 0.002 | 0.870 | 0.020 | 0.138 |

*Note.* Standardised betas (*β*) and *p* values are reported from regression models where each vascular risk factor and their age interaction are individually regressed onto volumetric MRI measures, covarying for sex, age, $age^2$, sex × age, sex × $age^2$, ethnicity and head position MRI confounds (volumetric data are also corrected for head size). GM grey matter volume; BMI = body mass index; WHR = waist:hip ratio.

*Supplementary Table 7.* Associations of aggregate vascular risk with tract-averaged white matter water diffusion parameters and subcortical volumes.

| | *Fractional Anisotropy* | | *Mean Diffusivity* | |
|---|---|---|---|---|
| ***Tracts*** | ***β*** | ***p*** | ***β*** | ***p*** |
| AR | 0.012 | 0.262 | 0.004 | 0.735 |
| ATR | **-0.037** | **0.001** | **0.082** | **<0.001** |
| CingG | **-0.047** | **<0.001** | **0.064** | **<0.001** |
| CingPH | 0.016 | 0.161 | 0.000 | 0.990 |
| CST | **0.052** | **<0.001** | **0.029** | **0.010** |
| FMaj | **-0.035** | **0.002** | **0.027** | **0.014** |
| FMin | **-0.062** | **<0.001** | **0.051** | **<0.001** |
| ILF | **-0.041** | **<0.001** | **0.050** | **<0.001** |
| IFOF | **-0.037** | **0.001** | **0.067** | **<0.001** |
| MCP | **0.065** | **<0.001** | **0.105** | **<0.001** |
| ML | 0.008 | 0.501 | **-0.060** | **<0.001** |
| PTR | **-0.094** | **<0.001** | **0.090** | **<0.001** |
| SLF | **-0.056** | **<0.001** | **0.074** | **<0.001** |
| STR | -0.005 | 0.656 | **0.078** | **<0.001** |
| Unc | -0.015 | 0.162 | **0.045** | **<0.001** |

| | *Volume* | |
|---|---|---|
| ***Subcortex*** | ***β*** | ***p*** |
| Accumbens | **-0.087** | **<0.001** |
| Amygdala | -0.006 | 0.576 |
| Caudate | **-0.046** | **<0.001** |
| Hippocampus | **-0.059** | **<0.001** |
| Pallidum | **-0.077** | **<0.001** |
| Putamen | **-0.058** | **<0.001** |
| Thalamus | **-0.073** | **<0.001** |

*Note.* Standardised betas (*β*) and *p* values reported, associations that survive FDR correction in bold. AR: acoustic radiation, ATR: anterior thalamic radiation, IFOF: inferior fronto-occipital fasciculus, ILF: inferior longitudinal fasciculus, MCP: middle cerebellar peduncle, PH: parahippocampal portion, PTR: posterior thalamic radiation, SLF: superior longitudinal fasciculus, STR: superior thalamic radiation.

*Supplementary Table 8.* Associations of latent vascular risk with tract-averaged white matter water diffusion parameters and subcortical volumes.

| | *Fractional Anisotropy* | | *Mean Diffusivity* | |
|---|---|---|---|---|
| ***Tracts*** | ***β*** | ***p*** | ***β*** | ***p*** |
| AR | 0.023 | 0.071 | 0.000 | 0.996 |
| ATR | **-0.033** | **0.010** | **0.088** | **<0.001** |
| CingG | **-0.042** | **0.001** | **0.072** | **<0.001** |
| CingPH | **0.035** | **0.007** | -0.023 | 0.078 |
| CST | **0.093** | **<0.001** | 0.012 | 0.334 |
| FMaj | -0.026 | 0.045 | 0.021 | 0.090 |
| FMin | **-0.064** | **<0.001** | **0.048** | **<0.001** |
| ILF | **-0.035** | **0.006** | **0.051** | **<0.001** |
| IFOF | **-0.031** | **0.016** | **0.072** | **<0.001** |
| MCP | **0.107** | **<0.001** | **0.133** | **<0.001** |
| ML | **0.028** | **0.031** | **-0.097** | **<0.001** |
| PTR | **-0.088** | **<0.001** | **0.080** | **<0.001** |
| SLF | **-0.053** | **<0.001** | **0.073** | **<0.001** |
| STR | 0.008 | 0.517 | **0.076** | **<0.001** |
| Unc | -0.011 | 0.398 | **0.055** | **<0.001** |

| | *Volume* | |
|---|---|---|
| ***Subcortex*** | ***β*** | ***p*** |
| Accumbens | **-0.096** | **<0.001** |
| Amygdala | -0.013 | 0.267 |
| Caudate | **-0.071** | **<0.001** |
| Hippocampus | **-0.064** | **<0.001** |
| Pallidum | **-0.107** | **<0.001** |
| Putamen | **-0.079** | **<0.001** |
| Thalamus | **-0.086** | **<0.001** |

*Note.* Standardised betas (*β*) and *p* values reported, associations that survive FDR correction in bold. AR: acoustic radiation, ATR: anterior thalamic radiation, IFOF: inferior fronto-occipital fasciculus, ILF: inferior longitudinal fasciculus, MCP: middle cerebellar peduncle, PH: parahippocampal portion, PTR: posterior thalamic radiation, SLF: superior longitudinal fasciculus, STR: superior thalamic radiation.

*Supplementary Table 9.* Associations between individually-modelled vascular risk factors and white matter tract fractional anisotropy.

| **Vascular Risk** | **AR** | **ATR** | **CingG** | **CingP** | **CST** | **FMaj** | **FMin** | **IFOF** | **ILF** | **MCP** | **ML** | **PTR** | **SLF** | **STR** | **Unc** |
|---|---|---|---|---|---|---|---|---|---|---|---|---|---|---|---|
| Pack Years | -0.004 | **-0.035** | **-0.038** | -0.009 | 0.009 | -0.018 | **-0.041** | **-0.034** | **-0.041** | 0.022 | 0.007 | **-0.062** | **-0.030** | **-0.035** | -0.021 |
| Hypertension | **-0.030** | **-0.085** | **-0.045** | -0.021 | 0.003 | **-0.040** | **-0.093** | **-0.073** | **-0.064** | 0.006 | -0.024 | **-0.087** | **-0.087** | **-0.050** | **-0.052** |
| Pulse Pressure | **-0.034** | **-0.032** | -0.021 | 0.025 | 0.003 | **-0.028** | **-0.034** | **-0.040** | **-0.042** | 0.000 | 0.017 | -0.015 | **-0.050** | -0.025 | **-0.026** |
| Diabetes | -0.010 | **-0.031** | **-0.033** | -0.001 | **0.026** | -0.021 | **-0.042** | **-0.034** | **-0.037** | 0.005 | -0.003 | **-0.057** | **-0.037** | -0.004 | **-0.024** |
| HiCholesterol | -0.008 | -0.005 | -0.015 | **0.027** | 0.008 | -0.020 | 0.003 | -0.002 | -0.005 | 0.022 | -0.001 | -0.021 | -0.004 | -0.009 | -0.005 |
| BMI | **0.041** | 0.007 | -0.017 | **0.026** | **0.083** | -0.006 | **-0.027** | 0.010 | -0.002 | **0.116** | 0.022 | **-0.043** | -0.022 | **0.028** | 0.018 |
| WHR | **0.033** | 0.000 | -0.019 | **0.045** | **0.088** | -0.005 | -0.022 | -0.004 | -0.004 | **0.087** | **0.040** | **-0.049** | -0.011 | 0.029 | 0.008 |

*Note*. Standardised $\beta$s reported. Bold type indicates FDR significant ($q < 0.05$). AR: acoustic radiation, ATR: anterior thalamic radiation, IFOF: inferior fronto-occipital fasciculus, ILF: inferior longitudinal fasciculus, MCP: middle cerebellar peduncle, PH: parahippocampal portion, PTR: posterior thalamic radiation, SLF: superior longitudinal fasciculus, STR: superior thalamic radiation, BMI: body mass index, WHR: waist:hip ratio.

*Supplementary Table 10.* Associations between individually-modelled vascular risk factors and white matter tract mean diffusivity.

| **Vascular Risk** | **AR** | **ATR** | **CingG** | **CingP** | **CST** | **FMaj** | **FMin** | **IFOF** | **ILF** | **MCP** | **ML** | **PTR** | **SLF** | **STR** | **Unc** |
|---|---|---|---|---|---|---|---|---|---|---|---|---|---|---|---|
| Pack Years | -0.005 | **0.041** | **0.026** | 0.014 | 0.019 | 0.013 | **0.033** | **0.045** | **0.041** | **0.062** | -0.005 | **0.062** | **0.037** | **0.055** | **0.023** |
| Hypertension | **0.046** | **0.104** | **0.068** | 0.007 | **0.054** | **0.025** | **0.083** | **0.097** | **0.080** | **0.041** | 0.000 | **0.076** | **0.106** | **0.089** | **0.067** |
| Pulse Pressure | **0.057** | **0.068** | **0.053** | -0.023 | **0.033** | **0.037** | **0.062** | **0.075** | **0.068** | 0.011 | -0.023 | **0.044** | **0.067** | **0.066** | **0.057** |
| Diabetes | 0.017 | **0.056** | **0.023** | 0.020 | **0.041** | 0.008 | **0.027** | **0.035** | **0.029** | **0.085** | -0.005 | **0.044** | **0.036** | **0.059** | **0.043** |
| HiCholesterol | 0.013 | 0.014 | 0.014 | -0.020 | 0.016 | **0.026** | -0.002 | 0.008 | 0.009 | 0.011 | **-0.030** | **0.023** | 0.015 | **0.023** | 0.005 |
| BMI | **-0.035** | **0.022** | **0.031** | **-0.034** | **-0.041** | 0.000 | 0.008 | 0.014 | -0.003 | **0.093** | **-0.111** | **0.023** | 0.015 | 0.009 | 0.003 |
| WHR | -0.010 | **0.055** | **0.051** | -0.022 | 0.005 | 0.013 | 0.017 | **0.040** | **0.027** | **0.099** | **-0.076** | **0.055** | **0.038** | **0.043** | **0.035** |

*Note*. Standardised $\beta$s reported. Bold type indicates FDR significant ($q < 0.05$). AR: acoustic radiation, ATR: anterior thalamic radiation, IFOF: inferior fronto-occipital fasciculus, ILF: inferior longitudinal fasciculus, MCP: middle cerebellar peduncle, PH: parahippocampal portion, PTR: posterior thalamic radiation, SLF: superior longitudinal fasciculus, STR: superior thalamic radiation, BMI: body mass index, WHR: waist:hip ratio.

*Supplementary Table 11.* Associations between individually-modelled vascular risk factors on subcortical volumes.

| Risk Factors | Accumbens | Amygdala | Caudate | Hippocampus | Pallidum | Putamen | Thalamus |
|---|---|---|---|---|---|---|---|
| Pack Years | **-0.047 (<0.001)** | -0.005 (0.590) | **-0.024 (0.006)** | **-0.032 (<0.001)** | **-0.046 (<0.001)** | **-0.044 (<0.001)** | **-0.054 (<0.001)** |
| Hypertension | **-0.061 (<0.001)** | **-0.041 (<0.001)** | -0.007 (0.394) | **-0.045 (<0.001)** | **-0.032 (<0.001)** | **-0.031 (<0.001)** | **-0.036 (<0.001)** |
| Pulse Pressure | 0.013 (0.190) | 0.002 (0.875) | -0.001 (0.937) | -0.010 (0.296) | -0.013 (0.196) | 0.005 (0.545) | -0.001 (0.916) |
| Diabetes | **-0.059 (<0.001)** | -0.011 (0.249) | **-0.030 (<0.001)** | **-0.042 (<0.001)** | **-0.064 (<0.001)** | **-0.045 (<0.001)** | **-0.051 (<0.001)** |
| HiCholesterol | **-0.028 (0.002)** | -0.011 (0.233) | -0.001 (0.871) | -0.017 (0.062) | -0.005 (0.566) | -0.008 (0.333) | **-0.019 (0.006)** |
| BMI | **-0.060 (<0.001)** | 0.018 (0.058) | **-0.052 (<0.001)** | -0.020 (0.032) | **-0.075 (<0.001)** | **-0.062 (<0.001)** | **-0.049 (<0.001)** |
| WHR | **-0.070 (<0.001)** | -0.019 (0.136) | **-0.072 (<0.001)** | **-0.062 (<0.001)** | **-0.085 (<0.001)** | **-0.055 (<0.001)** | **-0.070 (<0.001)** |

*Note.* Standardised betas ($\beta$) and *p* values are reported from regression models where each individual vascular risk factor is separately regressed onto subcortical volumes corrected for age, sex, ethcnicity, head size and scanner head position confounds. Bold type denotes FDR q-value < 0.05. BMI = body mass index; WHR = waist:hip ratio.

*Supplementary Table 11.* Associations between simultaneously-modelled vascular risk factors and white matter tract fractional anisotropy.

| Vascular Risk | AR | ATR | CingG | CingP | CST | FMaj | FMin | IFOF | ILF | MCP | ML | PTR | SLF | STR | Unc |
|---|---|---|---|---|---|---|---|---|---|---|---|---|---|---|---|
| Pack Years | -0.010 | **-0.030** | **-0.032** | -0.018 | -0.004 | -0.018 | **-0.037** | **-0.035** | **-0.041** | 0.010 | -0.001 | **-0.052** | -0.026 | **-0.039** | -0.021 |
| Hypertension | **-0.033** | **-0.089** | **-0.044** | **-0.031** | -0.016 | **-0.034** | **-0.091** | **-0.072** | **-0.061** | -0.016 | **-0.031** | **-0.079** | **-0.083** | **-0.056** | **-0.053** |
| Pulse Pressure | **-0.033** | -0.019 | -0.013 | 0.028 | -0.001 | -0.022 | -0.018 | **-0.031** | **-0.033** | -0.004 | 0.019 | -0.002 | **-0.037** | -0.021 | -0.019 |
| Diabetes | -0.007 | -0.017 | -0.019 | -0.003 | 0.015 | -0.012 | -0.024 | -0.023 | -0.025 | -0.013 | -0.003 | **-0.035** | -0.019 | 0.005 | -0.017 |
| HiCholesterol | -0.010 | 0.007 | -0.005 | 0.027 | -0.001 | -0.015 | 0.018 | 0.009 | 0.003 | 0.015 | 0.000 | -0.003 | 0.008 | -0.004 | 0.000 |
| BMI | **0.047** | 0.029 | 0.000 | 0.018 | **0.070** | 0.005 | -0.003 | **0.034** | 0.018 | **0.113** | 0.013 | -0.010 | -0.003 | **0.036** | **0.031** |
| WHR | 0.012 | 0.003 | 0.000 | 0.038 | **0.041** | 0.003 | 0.005 | -0.003 | 0.007 | 0.010 | 0.037 | -0.015 | 0.010 | 0.023 | 0.002 |

*Note*. Standardised $\beta$s reported. Bold type indicates FDR significant ($q < 0.05$). AR: acoustic radiation, ATR: anterior thalamic radiation, IFOF: inferior fronto-occipital fasciculus, ILF: inferior longitudinal fasciculus, MCP: middle cerebellar peduncle, PH: parahippocampal portion, PTR: posterior thalamic radiation, SLF: superior longitudinal fasciculus, STR: superior thalamic radiation, BMI: body mass index, WHR: waist:hip ratio.

*Supplementary Table 12.* Associations between simultaneously-modelled vascular risk factors and white matter tract mean diffusivity.

| Vascular Risk | AR | ATR | CingG | CingP | CST | FMaj | FMin | IFOF | ILF | MCP | ML | PTR | SLF | STR | Unc |
|---|---|---|---|---|---|---|---|---|---|---|---|---|---|---|---|
| Pack Years | -0.001 | **0.026** | 0.018 | 0.018 | 0.013 | 0.014 | **0.031** | **0.040** | **0.039** | **0.047** | 0.011 | **0.056** | **0.032** | **0.047** | 0.016 |
| Hypertension | **0.045** | **0.094** | **0.059** | 0.011 | **0.055** | 0.018 | **0.078** | **0.088** | **0.074** | 0.020 | 0.022 | **0.065** | **0.098** | **0.077** | **0.061** |
| Pulse Pressure | **0.054** | **0.051** | **0.042** | -0.026 | 0.025 | **0.035** | **0.049** | **0.061** | **0.057** | -0.003 | -0.020 | **0.032** | **0.052** | **0.052** | **0.047** |
| Diabetes | 0.013 | **0.036** | 0.010 | 0.025 | **0.041** | 0.001 | 0.013 | 0.019 | 0.017 | **0.067** | 0.013 | **0.026** | 0.017 | **0.042** | **0.034** |
| HiCholesterol | 0.011 | -0.002 | 0.003 | -0.019 | 0.009 | 0.025 | -0.012 | -0.006 | -0.002 | -0.006 | -0.022 | 0.008 | 0.000 | 0.009 | -0.005 |
| BMI | **-0.054** | -0.021 | 0.002 | **-0.035** | **-0.072** | -0.014 | -0.016 | -0.024 | **-0.036** | **0.061** | **-0.113** | -0.018 | -0.022 | **-0.033** | -0.027 |
| WHR | 0.011 | **0.038** | 0.031 | -0.004 | 0.032 | 0.012 | 0.003 | 0.027 | 0.025 | 0.036 | -0.005 | **0.039** | 0.025 | 0.030 | 0.031 |

*Note*. Standardised $\beta$s reported. Bold type indicates FDR significant ($q < 0.05$). AR: acoustic radiation, ATR: anterior thalamic radiation, IFOF: inferior fronto-occipital fasciculus, ILF: inferior longitudinal fasciculus, MCP: middle cerebellar peduncle, PH: parahippocampal portion, PTR: posterior thalamic radiation, SLF: superior longitudinal fasciculus, STR: superior thalamic radiation, BMI: body mass index, WHR: waist:hip ratio.

*Supplementary Table 13.* Associations between simultaneously-modelled vascular risk factors on subcortical volumes.

| **Risk Factors** | **Accumbens** | **Amygdala** | **Caudate** | **Hippocampus** | **Pallidum** | **Putamen** | **Thalamus** |
|---|---|---|---|---|---|---|---|
| Pack Years | **-0.029 (0.001)** | -0.002 (0.827) | -0.016 (0.064) | **-0.024 (0.011)** | **-0.032 (<0.001)** | **-0.035 (<0.001)** | **-0.042 (<0.001)** |
| Hypertension | **-0.049 (<0.001)** | **-0.044 (<0.001)** | 0.005 (0.553) | **-0.037 (<0.001)** | -0.014 (0.124) | **-0.021 (0.011)** | **-0.022 (0.003)** |
| Pulse Pressure | **0.026 (0.009)** | 0.006 (0.547) | 0.003 (0.716) | -0.003 (0.757) | -0.005 (0.614) | 0.015 (0.094) | 0.008 (0.312) |
| Diabetes | **-0.041 (<0.001)** | -0.005 (0.643) | **-0.022 (0.014)** | **-0.032 (<0.001)** | **-0.046 (<0.001)** | **-0.033 (<0.001)** | **-0.035 (<0.001)** |
| HiCholesterol | -0.012 (0.193) | -0.005 (0.605) | 0.005 (0.582) | -0.007 (0.436) | 0.008 (0.375) | 0.004 (0.639) | -0.008 (0.279) |
| BMI | **-0.033 (0.002)** | **0.041 (<0.001)** | **-0.031 (0.003)** | 0.015 (0.162) | **-0.048 (<0.001)** | **-0.047 (<0.001)** | **-0.023 (0.007)** |
| WHR | -0.023 (0.106) | **-0.040 (0.009)** | **-0.046 (<0.001)** | **-0.057 (<0.001)** | **-0.034 (0.014)** | -0.004 (0.775) | **-0.036 (0.001)** |

*Note.* Standardised betas ($\beta$) and $p$ values are reported from regression models where each individual vascular risk factor is regressed onto subcortical volumes corrected for age, sex, ethcnicity, head size and scanner head position confounds. Bold type denotes FDR q-value < 0.05. BMI = body mass index; WHR = waist:hip ratio.

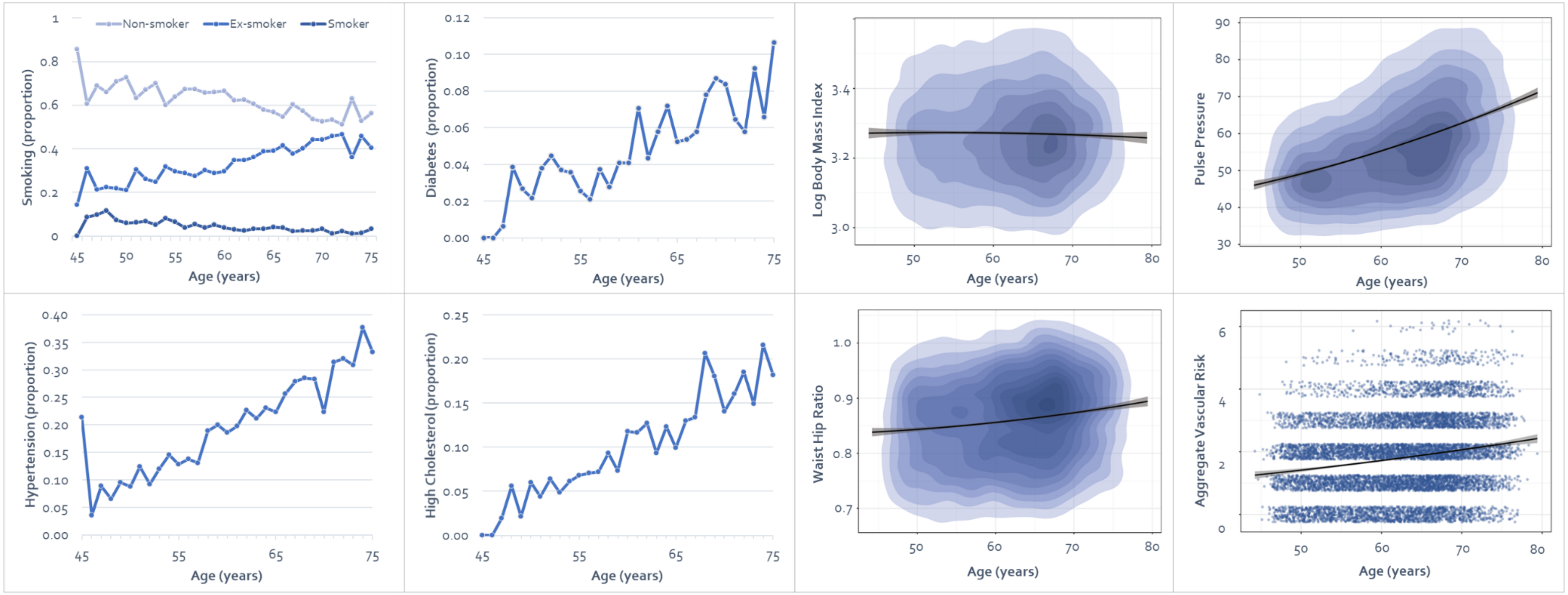


*Supplementary Figure 1*. Differences in vascular risk by age in UK Biobank. From top left to bottom right: smoking status, diabetes, body mass index, pulse pressure, hypertension, hypercholesterolaemia, waist:hip ratio and aggregate vascular risk. The latter contains some vertical jitter for visualisation purposes. Regression lines (which allow a quadratic component) with shaded 95% CIs are shown in plots on the right hand side. Results reported in Table 1.

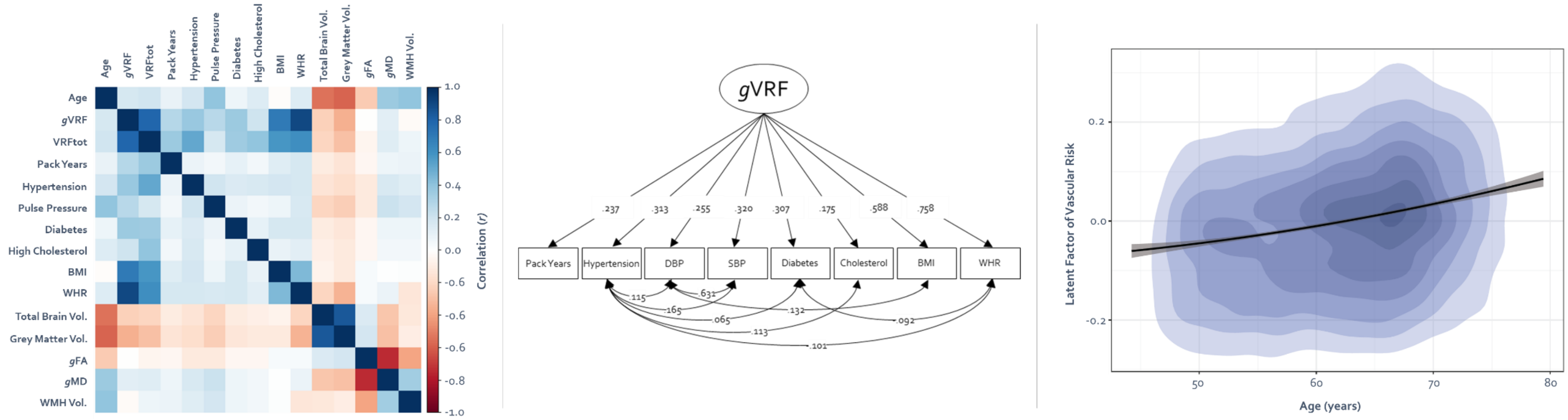


*Supplementary Figure 2.* Modelling common variance among vascular risk factors. **Left panel**: a correlogram of study variables illustrates the positive associations among vascular risk factors (see Supplementary Table 4 for correlation matrix). **Middle panel**: a path diagram illustrating the latent factor of vascular risk. **Right panel**: association between the latent factor of vascular risk and age with egression lines and shaded 95% CI.

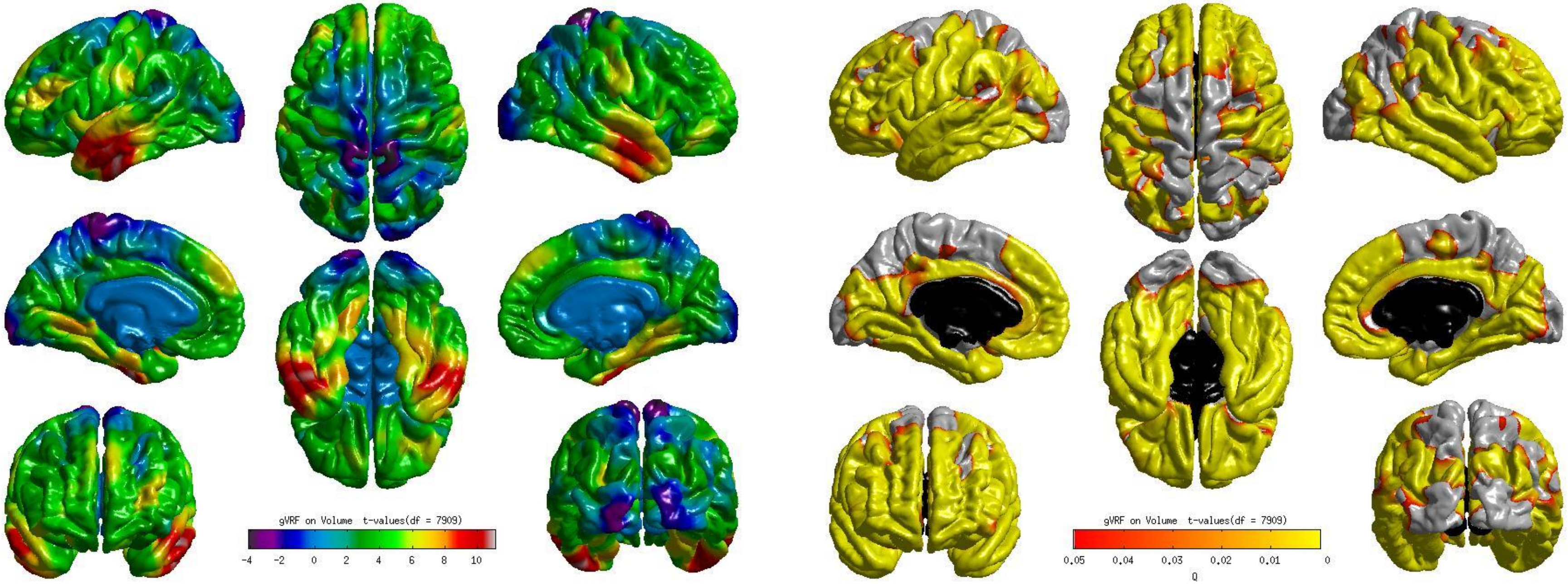


*Supplementary Figure 3.* Associations (**left**: t-maps and **right**: FDR-corrected *q*-maps) between a latent factor of vascular risk and cortical volume.

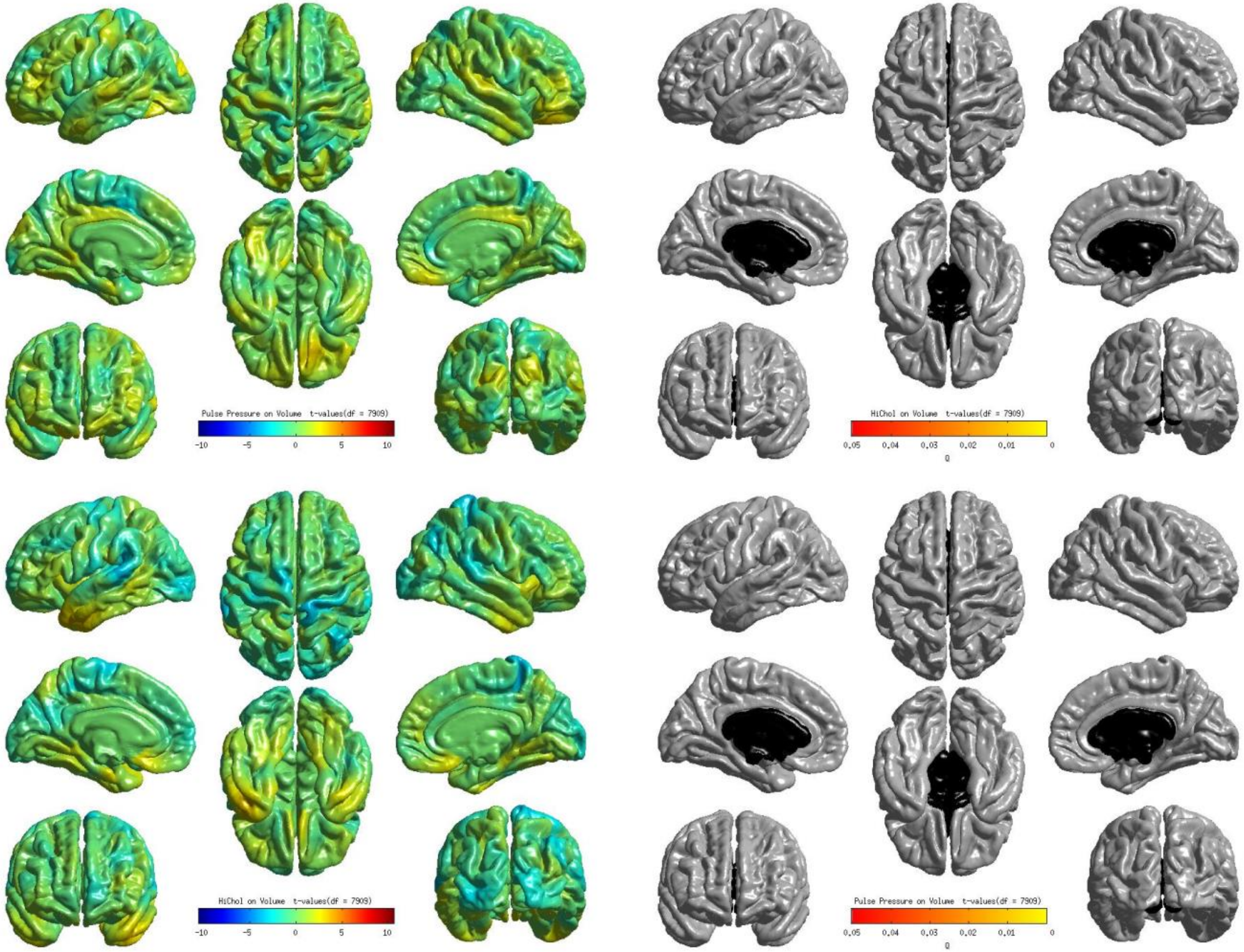


*Supplementary Figure 4.* Associations (**left:** t-maps and **right:** FDR-corrected *q*-maps) between cortical volume and pulse pressure (top row) and hypercholesterolaemia (bottom row), modelled individually. See Figure 5 for FDR-corrected significant results.

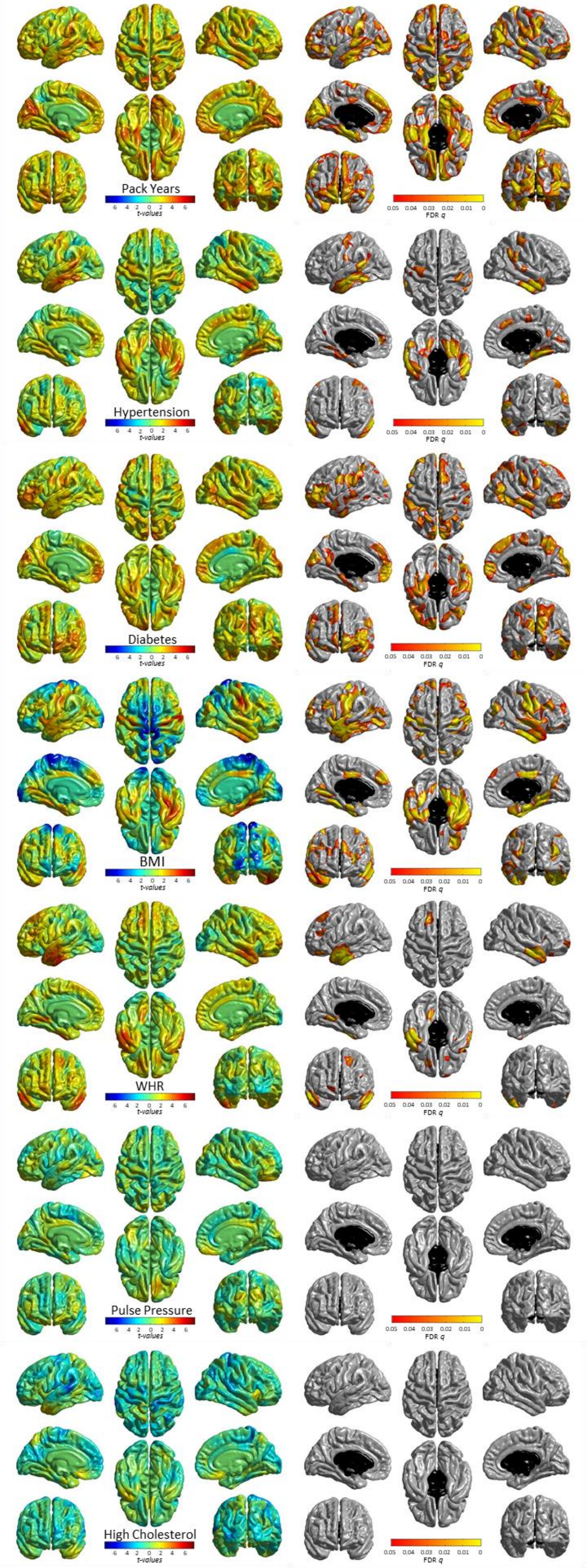


*Supplementary Figure 5.* Significant associations (**left:** t-maps and **right:** FDR-corrected *q*-values) between cortical volume and vascular risk factors (modelled simultaneously, alongside age, sex, ethnicity, head size and scanner head position confounds).

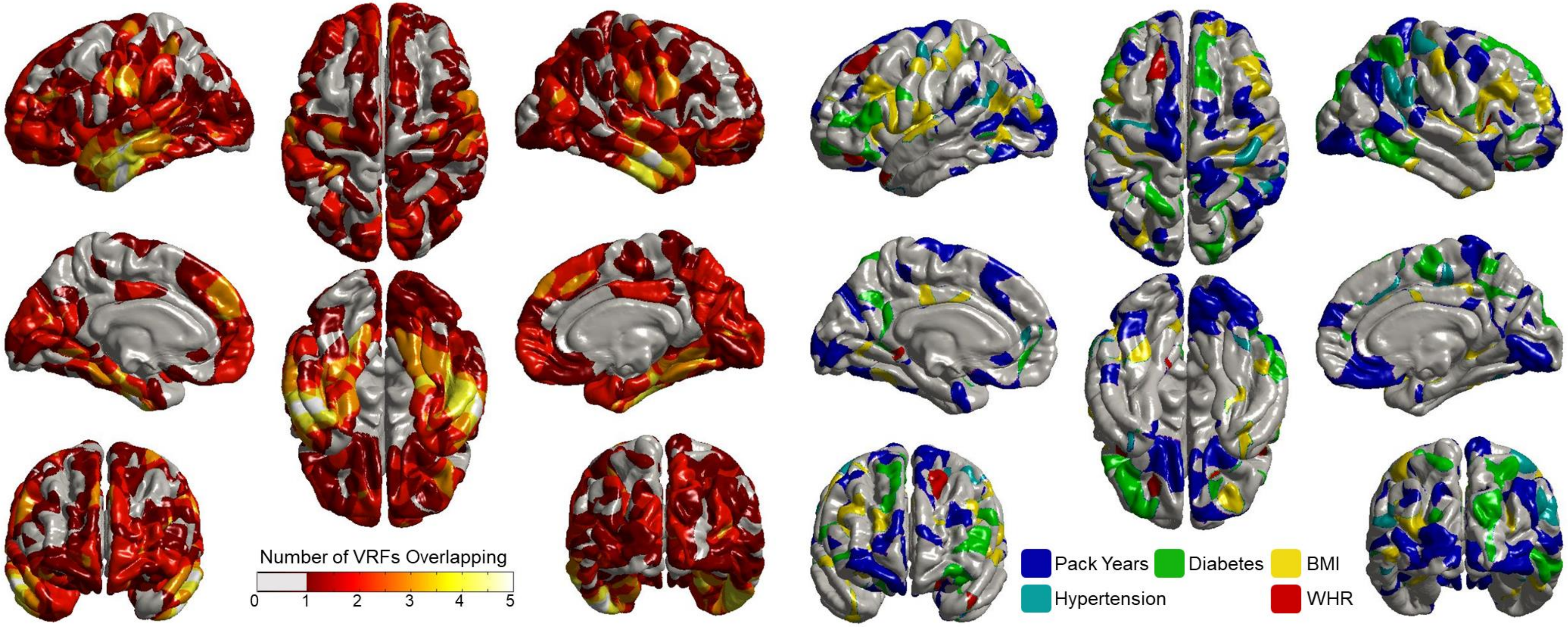


*Supplementary Figure 6.* Conjunction plot (**left**) shows the cortical loci that exhibit unique associations with simultaneously-modelled vascular risk factors (VRFs). The conditional analysis **(right**) decomposes the non-overlapping areas (dark red; left), assigning each locus to its respective VRF, i.e. areas that are only affected by one VRF. The corresponding t-maps and FDR-corrected results for the unique contributions of each VRF are shown in Supplementary Figure 5.
